# Kinase-independent inhibition of cyclophosphamide-induced pathways protects the ovarian reserve and prolongs fertility

**DOI:** 10.1101/526103

**Authors:** Giovanna Bellusci, Valentina Iannizzotto, Sarah Ciccone, Luca Mattiello, Emiliano Maiani, Valentina Villani, Marc Diederich, Stefania Gonfloni

**Affiliations:** Department of Biology, University of Rome Tor Vergata, via della Ricerca Scientifica, 00133 Rome, Italy; College of Pharmacy, Seoul National University, 1 Gwanak-ro, Gwanak-gu, Seoul 08826, Republic of Korea

## Abstract

Premature ovarian failure and infertility are adverse effects of cancer therapies. The mechanism underlying chemotherapy-mediated depletion of the ovarian reserve remains unclear. Here, we aim to identify the signaling pathways involved in the loss of the ovarian reserve to prevent the damaging effects of chemotherapy. We evaluated the effects of cyclophosphamide, one of the most damaging chemotherapeutic drugs, against follicle reserve. *In vivo* studies showed that the cyclophosphamide-induced loss of ovarian reserve occurred through a sequential mechanism. Cyclophosphamide exposure induced the activation of both DNAPK-γH2AX-checkpoint kinase 2 (CHK2)-p53/TAp63*α* isoform and protein kinase B (AKT)-forkhead box O3 (FOXO3) signaling axes in the nucleus of oocytes. Concomitant administration of an allosteric ABL inhibitor and cyclophosphamide modulated both pathways while protecting the ovarian reserve from chemotherapy assaults. As a consequence, the fertility of the treated mice was prolonged. On the contrary, the administration of an allosteric ABL activator enhanced the lethal effects of cyclophosphamide while shortening mouse fertility. Therefore, kinase-independent inhibition may serve as an effective ovarian-protective strategy in women under chemotherapy.

## Introduction

Premature ovarian failure and infertility are frequent consequences of cancer therapy^1^. Chemotherapy triggers the degeneration of the ovarian reserve^2, 3^, a source of fertilizable eggs during the entire reproductive life of a female. Preserving the ovarian reserve during cancer treatment is a major concern for the maintenance of fertility and to ameliorate the quality of life of survivors. Studies in transgenic mouse models have revealed the role of several molecules in the maintenance of ovarian reserve following ionization radiation (IR) treatment. The alpha TAp63 isoform (TAp63*α*) was thought to be a key mediator of IR-induced ovarian reserve loss^4^. Lack of TAp63*α* expression in mouse oocytes promoted resistance to the lethal effects of IR^5^. IR-induced TAp63 activation depended on a conformational transition of the molecule^6^. The pro-apoptotic proteins PUMA and Noxa may act as downstream effectors of TAp63*α*^7^. In mice, checkpoint kinase 2 (CHK2) is essential for DNA damage surveillance in female meiosis^8^. Moreover, the lack of CHK2 expression affects TAp63 activation following IR treatment^8^. *In vitro* experiments in ovarian culture system showed that the transient administration of a CHK2 inhibitor II hydrate may preserve oocyte viability following IR treatment^9^. Another study demonstrated that CHK2 inhibitor II hydrate affected the activities of a broad spectrum of kinases and impacted the global repression of the DNA damage response (DDR) in cultured ovaries^10^, thereby raising some questions about its specificity. Tuppi *et al* reported that CHK2 and the executioner kinase CK1 are both involved in the TAp63*α*-mediated oocyte degeneration in cultured ovaries exposed to chemotherapy^11^. As cultured mouse ovaries exposed to chemotherapeutic drugs may not recapitulate the signaling pathways physiologically occurring in the ovary *in vivo*, we aimed to perform all our experiments under *in vivo* conditions. We evaluated the effects of cyclophosphamide, a chemotherapeutic agent commonly used for the treatment of pediatric cancer patients. Cyclophosphamide is transformed in the liver into active alkylating metabolites, which induce the formation of DNA adducts. How cyclophosphamide contributes to ovarian follicle depletion is yet incompletely understood and is debatable^12^. Meirow *et al* suggest that the mechanism underlying cyclophosphamide-induced oocyte loss comprised the accelerated activation of primordial follicle that results in a “burnout” effect^13^. However, chemotherapeutic drugs may directly damage the primordial follicle and induce apoptosis. The extent of ovarian reserve loss also depends on the type and dosage of the chemotherapeutic agent^14^. Understanding the molecular basis underlying the effect of chemotherapy on quiescent primordial follicles is therefore essential for the identification of an effective co-treatment that may simultaneously preserve fertility in women^15, 16^. Here, we investigated the *in vivo* consequences of different doses of cyclophosphamide on the ovarian reserve. We used a pre-pubertal mouse model to evaluate the effect of transient administration of small molecule kinase inhibitors to identify putative ferto-protective adjuvants^17^ to limit the damaging effects of chemotherapy.

## Results

### Cyclophosphamide triggers a dose-dependent loss of primordial follicles

We investigated the effects of cyclophosphamide on perinatal mouse ovaries. We injected mice (at postnatal day 7, P7) with different doses of cyclophosphamide (ranging from 50 to 200 mg/kg). One day after injection, ovaries were dissected and analyzed with TdT-mediated dUTP nick-end labelling (TUNEL), immunofluorescence (IF), and immunohistochemistry (IHC) assays. Cyclophosphamide administration induced a dose-dependent increase in the number of TUNEL-positive granulosa cells surrounding the growing follicles (Fig. 1A). Cell death in the ovarian follicle reserve was observed with IF staining for cleaved poly ADP ribose polymerase (PARP) (Fig. 1B). We found apoptotic oocytes in the groups treated with low doses of cyclophosphamide (50-75 mg/kg), while higher doses of cyclophosphamide (100-200 mg/kg) activated cleaved PARP in the granulosa cells of large growing follicles (see dashed yellow box in Fig. 1B). We also detected the phosphorylation of the histone variant H2AX at Ser139 residue, also named as γH2AX, an early hallmark of DDR, in the nucleus of cyclophosphamide-damaged oocytes (Fig. 1C). Temporary hair loss was reported a week later in cyclophosphamide-injected animals (Fig. 1D).

**Figure 1.**
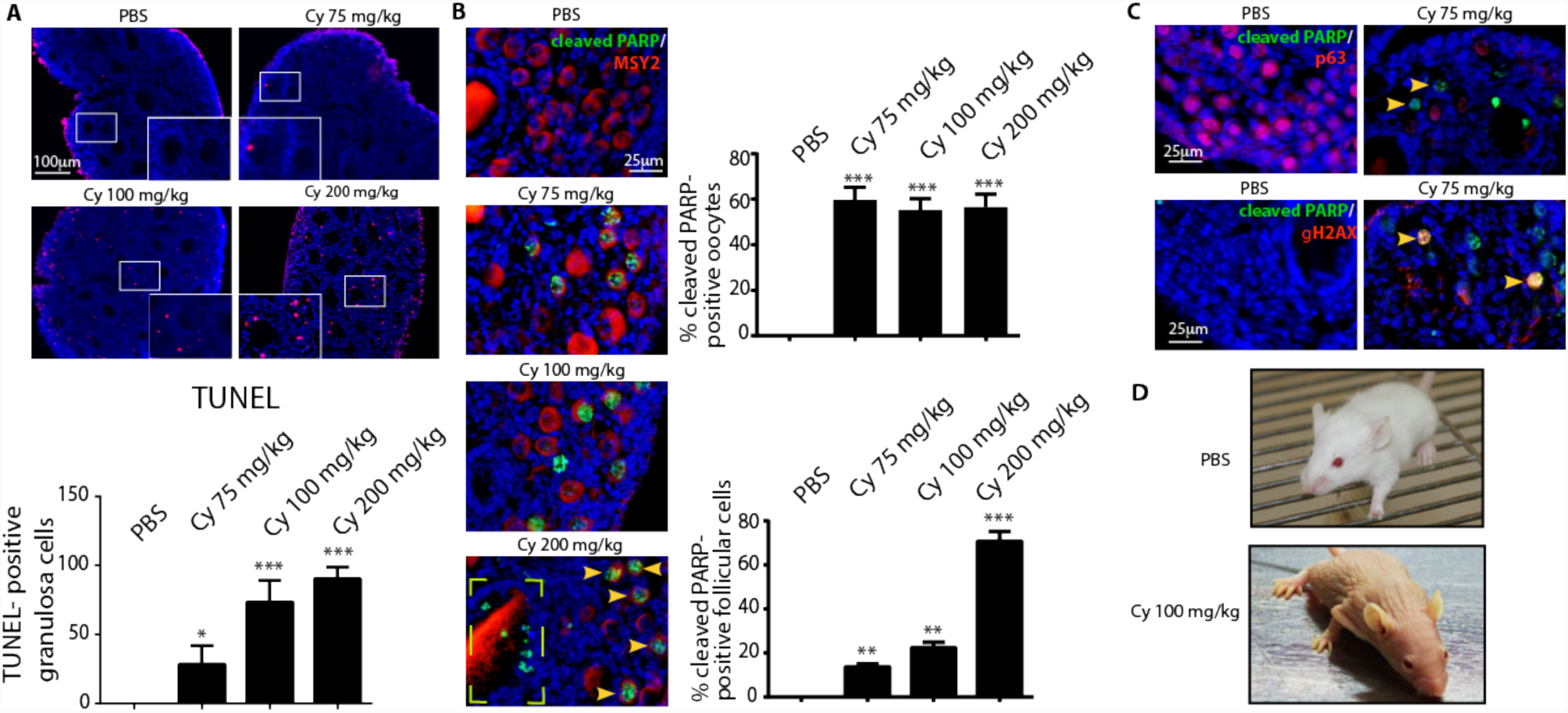
Cyclophosphamide triggers apoptosis in pre-pubertal mouse ovary in a dose-dependent manner. P7 mice were injected with vehicle (PBS) or increasing concentrations of cyclophosphamide (75, 100, and 200 mg/kg) and sacrificed within 16 h from injection. Ovarian sections were analyzed by in situ TdT-mediated dUTP nick-end labelling (TUNEL) assay. (A) A small area of each image was selected and zoomed in (magnification box). The graph shows the quantification of TUNEL-positive cells. Quantification of TUNEL-positive cells was performed by counting six different middle ovarian sections derived from three distinct ovaries. Follicle apoptosis was assessed by IF assay with two specific antibodies against cleaved PARP (green) and Msy2 (red), a cytoplasmic marker of germ cells. (B) Yellow arrows at the bottom indicate positive oocytes, while the dashed box illustrates PARP positive-granulosa cells from large growing follicle. Quantification of cleaved PARP-positive cells was performed by counting several (6 < x < 10) middle ovarian sections derived from three distinct ovaries. Scale bar magnification, 100 μm for TUNEL assay and 25 μm for IF assay. (C) Follicle apoptosis was assessed by IF assay with two specific antibodies against cleaved PARP (green) and p63 (red), a nuclear marker of germ cells. We also monitored γH2AX, a marker of DDR, in the reserve oocytes (yellow arrows). (D) Mouse images were obtained 7 days after cyclophosphamide injection. (A; B) Bar column represents mean ± s.d.; statistical significance was determined using one-way analysis of variance (ANOVA) (**P < 0.01; ***P < 0.001 as compared with PBS-treated group).

### DNA-PK is activated in the nucleus of reserve oocytes following cyclophosphamide treatment

To investigate the mechanism underlying cyclophosphamide-induced follicle death, we evaluated the activation of the apical DDR kinase (DNA-PK) with phospho-specific antibodies after 16 h of cyclophosphamide injection. The presence of apoptosis in the primordial/primary follicle population was consistent with the activation of DNA-PK in the nucleus of reserve oocytes (Fig. 2). Phosphorylation of the histone variant H2AX at Ser139 was observed in the nucleus of damaged oocytes. The majority of oocytes were positive for both DNA-PK and γH2AX expression at every dose of cyclophosphamide (Fig. 2). Confocal images of middle ovarian sections obtained from cyclophosphamide-injected mice showed that the oocytes with a strong signal for γH2AX or p-DNA-PK had lower expression of TAp63 (Fig. 2). Western blot analysis of the dissected ovaries revealed the cyclophosphamide-induced phosphorylation of both γH2AX and p53. In addition, cyclophosphamide induced TAp63 activation, as indicated by a mobility shift (*) that was absent in the control group (Fig. 2C).

**Figure 2.**
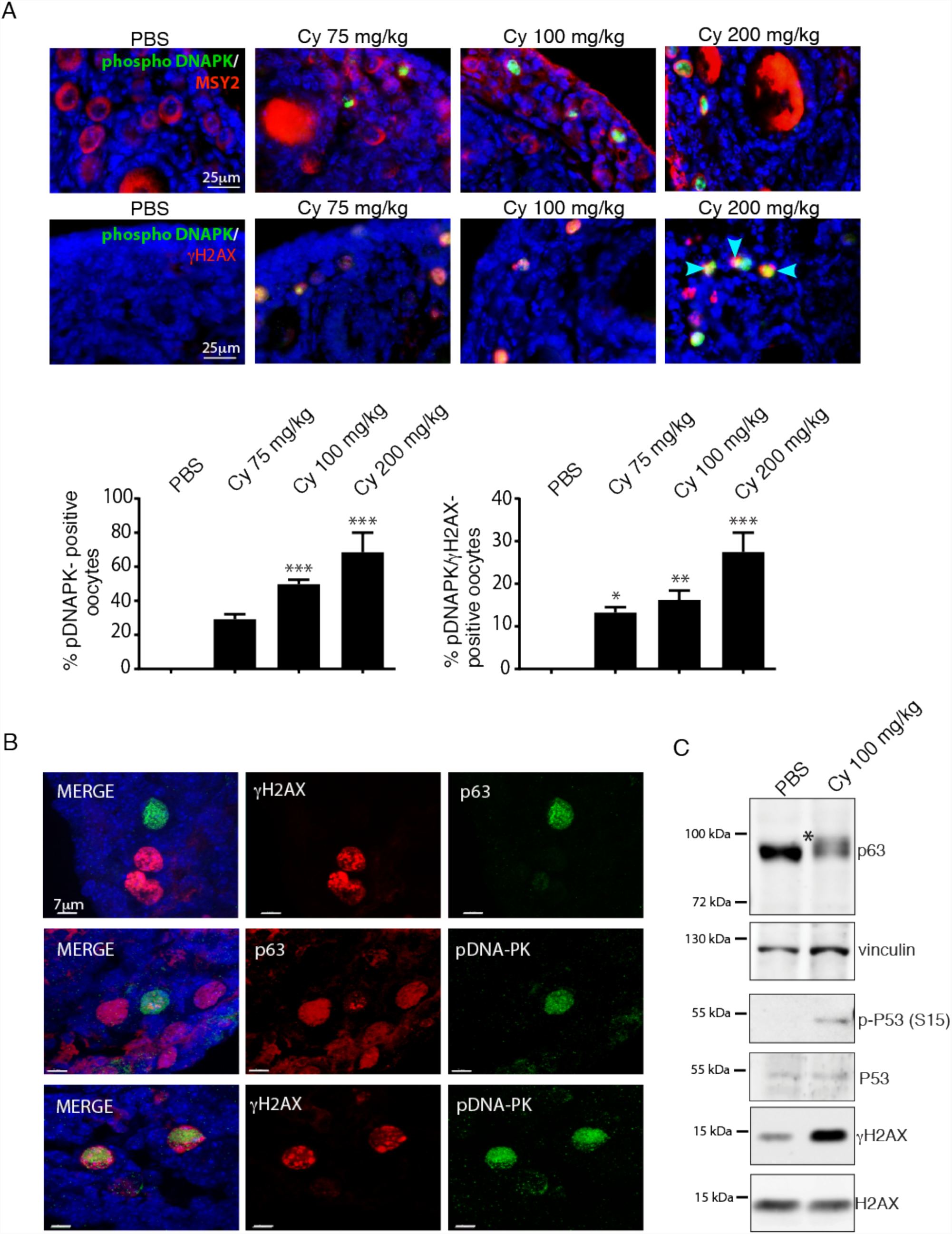
DNA-PK is activated in the nucleus of reserve oocytes following cyclophosphamide treatment. P7 mice were injected with vehicle (PBS) or increasing concentrations of cyclophosphamide (75, 100, and 200 mg/kg) and sacrificed within 16 h after injection. DNA-PK activation was evaluated with IF assay using a phospho-specific antibody (green) and Msy2 (red), a cytoplasmic antigen of germ cells (A). Blue arrows at the bottom indicate the oocytes positive for p-DNA-PK and γH2AX. Quantification was performed by counting several (6 < x < 8) middle ovarian sections derived from distinct ovaries. (B) Confocal images of middle ovarian sections from cyclophosphamide-injected mice indicated that the oocytes with high expression of p-DNA-PK or γH2AX showed reduced expression of TAp63. (C) Western blot analysis of ovarian lysates for female pups injected with 100 mg/kg of cyclophosphamide or PBS. Cyclophosphamide treatment induced a mobility shift of TAp63 (indicated with an asterisk [*]) that was absent in the control. Scale bar magnification, 25 μm for IF assay and 7 μm for confocal images. (A) Bar column represents mean ± s.d.; statistical significance was determined using one-way analysis of variance (ANOVA) (*P < 0.5; **P < 0.01; ***P < 0.001 as compared with PBS-treated group).

We observed activation of checkpoint kinase CHK2 and p53 in the nucleus of the damaged oocytes (Supplementary Fig. 1). Both CHK2 and p53 are involved in the removal of oocytes with unrepaired meiotic DNA double-strand breaks (DBSs)^8^. Thus, the cyclophosphamide-induced loss of ovarian reserve involves the activation of the DNA-PK-γH2AX-CHK2-p53/TAp63*α* signaling axes.

We investigated the mechanisms underlying the cyclophosphamide-induced follicle death following the activation of the protein kinase B (AKT) pathway using phospho-specific antibodies after 16 h treatment with cyclophosphamide (Supplementary Fig. 2). The AKT pathway is involved in primordial follicle activation through the regulation of the activity of the forkhead box O3 (FOXO3) transcription factor^18^. Upon phosphorylation, AKT may enter the nucleus and phosphorylate FOXO3, which eventually leaves the nucleus and enables follicular activation.

Following cyclophosphamide injection, p-AKT and p-FOXO3 were detected in the nucleus of reserve oocytes. We also observed expression of p-AKT and γH2AX in the reserve oocytes (Supplementary Fig. 2). Thus, the presence of the early marker of DDR was consistent with the activation of the AKT-FOXO3 pathway.

### GNF-2 prevents γH2AX phosphorylation in the nucleus of reserve oocytes

Previous studies have revealed the protective role of an ABL kinase inhibitor (imatinib) on ovarian reserve in cisplatin-induced degeneration^19, 20, 21, 22^. However, the mechanism underlying oocyte protection by imatinib remains unclear. As discussed in a recent paper^10^, cisplatin induced a response that activated the pathways different from those activated by IR. Cisplatin toxicity gradually accumulated from multiple sources (lipids, proteins, etc.) and induced integrated pathways of the two p53 homologs (TAp63*α* and TAp73*α*)^10^ in follicles. Here, we tested whether the allosteric ABL compounds affect the DDR induced by cyclophosphamide. In our experiments, we used an allosteric activator of ABL kinase^23^ (Supplementary Fig. 3) to validate the effect of ABL compounds *in vivo*. We treated female mice with cyclophosphamide in the presence of GNF-2. Concomitant administration of GNF-2 resulted in a significant reduction in the number of TUNEL-positive granulosa cells in a dose-dependent manner (Fig. 3). We failed to observe any apoptosis in reserve oocytes (Fig. 3). Co-treatment of cyclophosphamide and GNF-2 also affected DNA-PK activation and the consequent γH2AX phosphorylation in the nucleus of reserve oocytes (Fig. 3).

**Figure 3.**
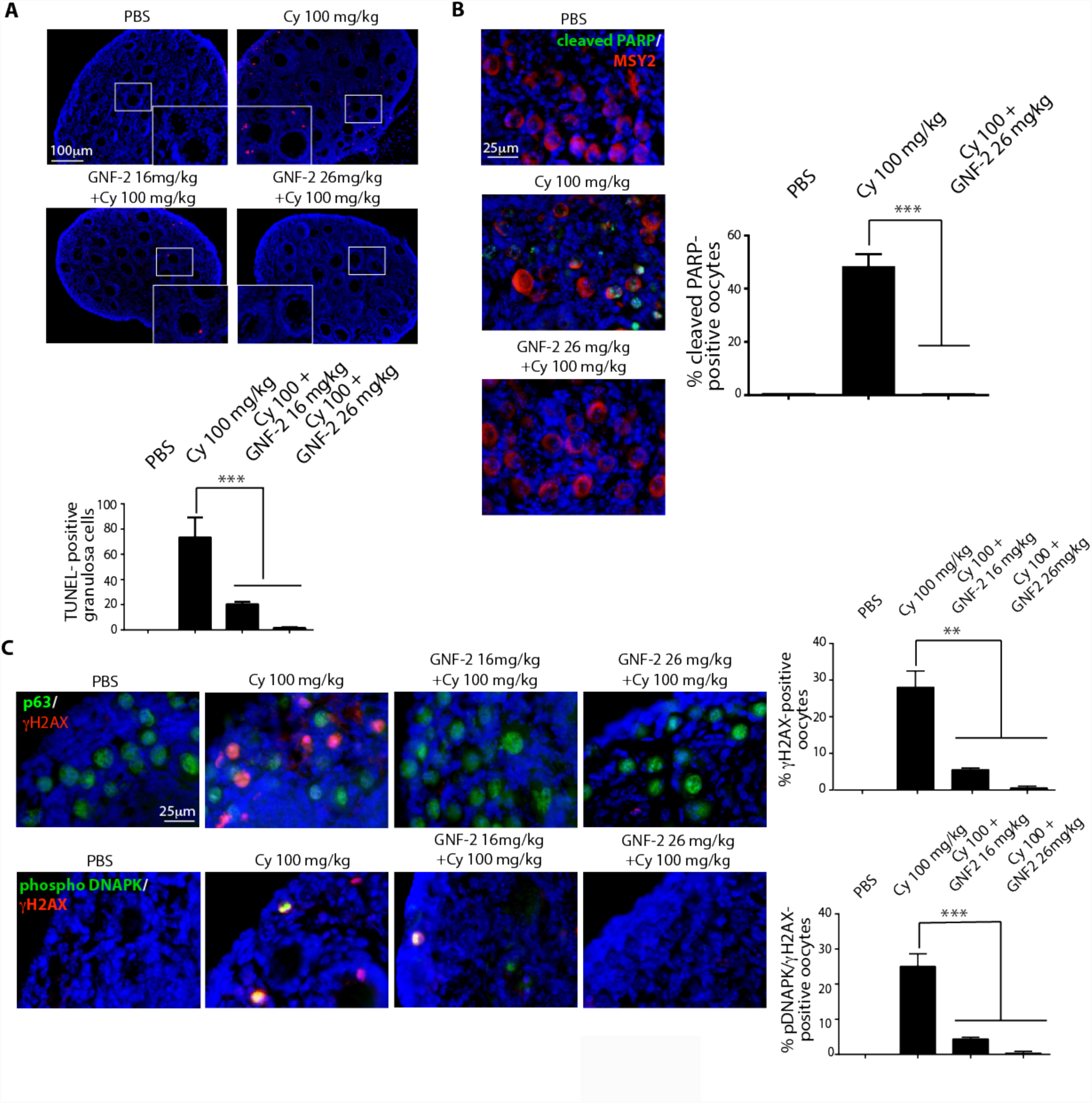
GNF-2 prevents the cyclophosphamide-induced DNA-PK-γH2AX signaling. P7 mice were injected with vehicle (PBS) or cyclophosphamide (100 mg/kg) with/without increasing concentrations of GNF-2 (16 and 26 mg/kg). Mice were sacrificed within 16 h after injection. (A) Ovarian sections were analyzed by in situ TdT-mediated dUTP nick-end labelling (TUNEL) assay. The graph shows the quantification of TUNEL-positive cells. Quantification of TUNEL-positive cells was performed by counting six different middle ovarian sections derived from three distinct ovaries. (B) Follicle reserve apoptosis was assessed by IF assay with two specific antibodies against cleaved PARP (green) and Msy2 (red), a cytoplasmic antigen of germ cells. Quantification of the cleaved PARP-positive cells was performed by counting several (6 < x < 8) middle ovarian sections derived from three distinct ovaries. (C) γH2AX and DNA-PK activation was investigated with IF assay using phospho-specific antibodies (red) and anti-p63 (green), a nuclear marker for germ cells. Quantification was performed by counting several (6 < x < 8) middle ovarian sections derived from three distinct ovaries. Scale bar magnification, 100 μm for TUNEL assay and 25 μm for IF assay. (A; B; C) Bar column represents mean ± s.d.; statistical significance was determined using one-way analysis of variance (ANOVA) (**P < 0.01; ***P < 0.001 as compared to 100 mg/kg cyclophosphamide).

On the contrary, the concomitant administration of DPH (an allosteric ABL activator) enhanced DDR activation (Fig. 4) and oocyte apoptosis induced by cyclophosphamide. IF assay results revealed the increase in the number of γH2AX-positive oocytes after cyclophosphamide treatment. Therefore, we investigated whether ATM phosphorylates H2AX in the damaged oocytes. IF assay results showed that ATM phosphorylation was coupled with a decrease in TAp63 expression in the nucleus of reserve oocytes, as reported for p-DNA-PK (Supplementary Fig. 4). The concomitant administration of GNF-2 also prevented ATM activation in the nucleus of reserve oocytes.

**Figure 4.**
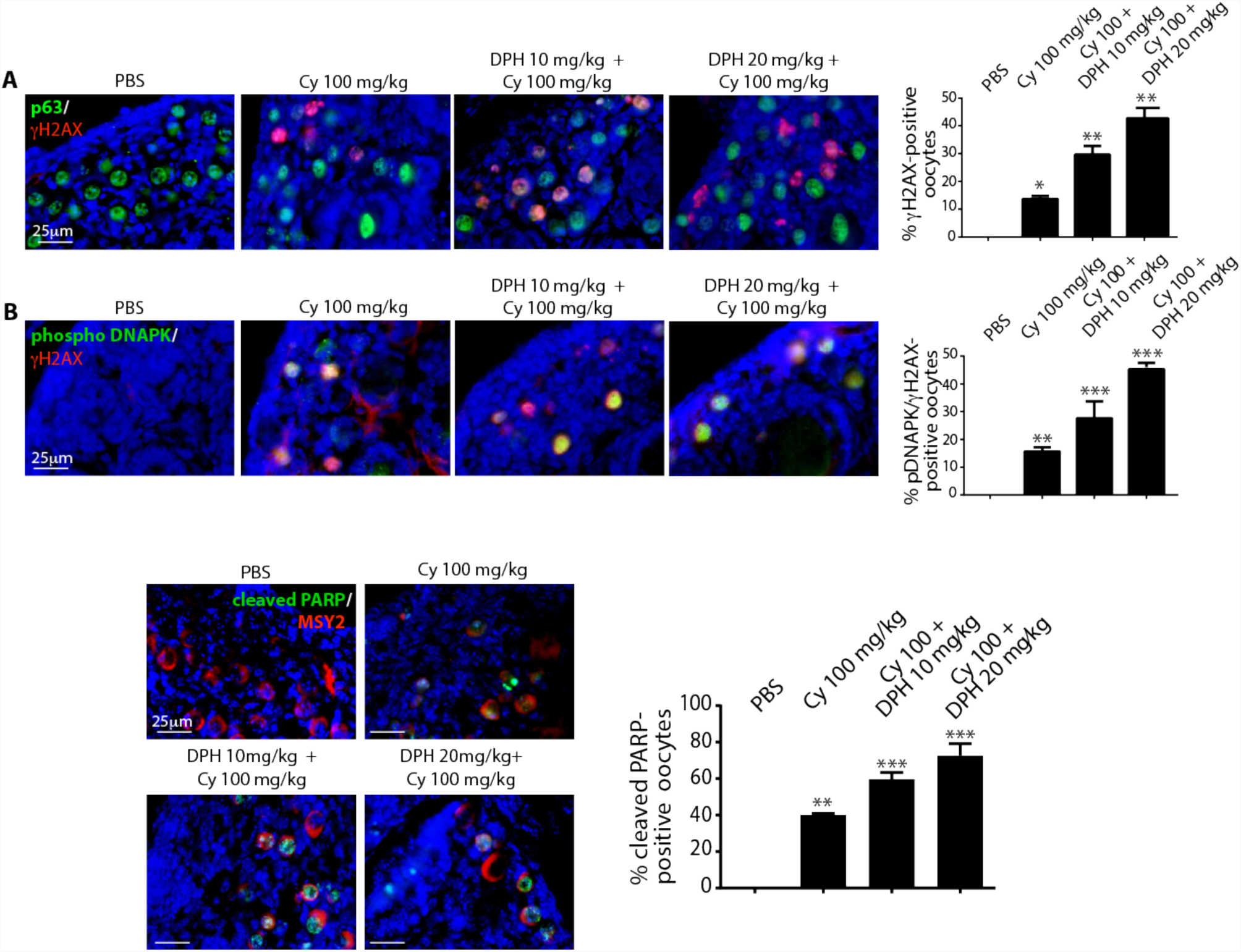
DPH enhances the DNA-PK-γH2AX signaling axis and oocyte apoptosis induced by cyclophosphamide. P7 mice were injected with vehicle (PBS) or cyclophosphamide (100 mg/kg) with/without increasing concentrations of DPH (10 and 20 mg/kg) and were sacrificed within 16 h from injection. (A, B) γH2AX and DNA-PK activation was evaluated with IF assay using phospho-specific antibodies (red), while p63 (green) was used as a nuclear marker for germ cells. Follicle reserve apoptosis was assessed by IF assay with two specific antibodies against cleaved PARP (green) and Msy2 (red), a cytoplasmic marker of germ cells. Quantification of cleaved PARP-positive cells was performed by counting several (6 < x < 8) middle ovarian sections derived from three distinct ovaries. Quantification was performed by counting several (6 < x < 8) middle ovarian sections derived from three distinct ovaries. Scale Bar magnification, 25 μm. Bar column represents mean ± s.d.; statistical significance was determined using one-way analysis of variance (ANOVA) (*P < 0.05; **P < 0.01; ***P < 0.001 as compared with PBS-treated group).

In conclusion, GNF-2 and cyclophosphamide co-treatment restricted apoptosis in ovarian reserve through the prevention of the activation of the DNA-PK(ATM)-γH2AX-CHK2-p53/TAp3 signaling axes. We also evaluated the effect of allosteric compounds on the activation of AKT-FOXO3 pathway. Co-treatment of cyclophosphamide with GNF-2 prevented the translocation of p-AKT in the nucleus of the oocyte, while DPH enhanced the activation of AKT and concomitant phosphorylation of FOXO3 (Supplementary Fig. 5) in the ovarian reserve assaulted by cyclophosphamide.

**Figure 5.**
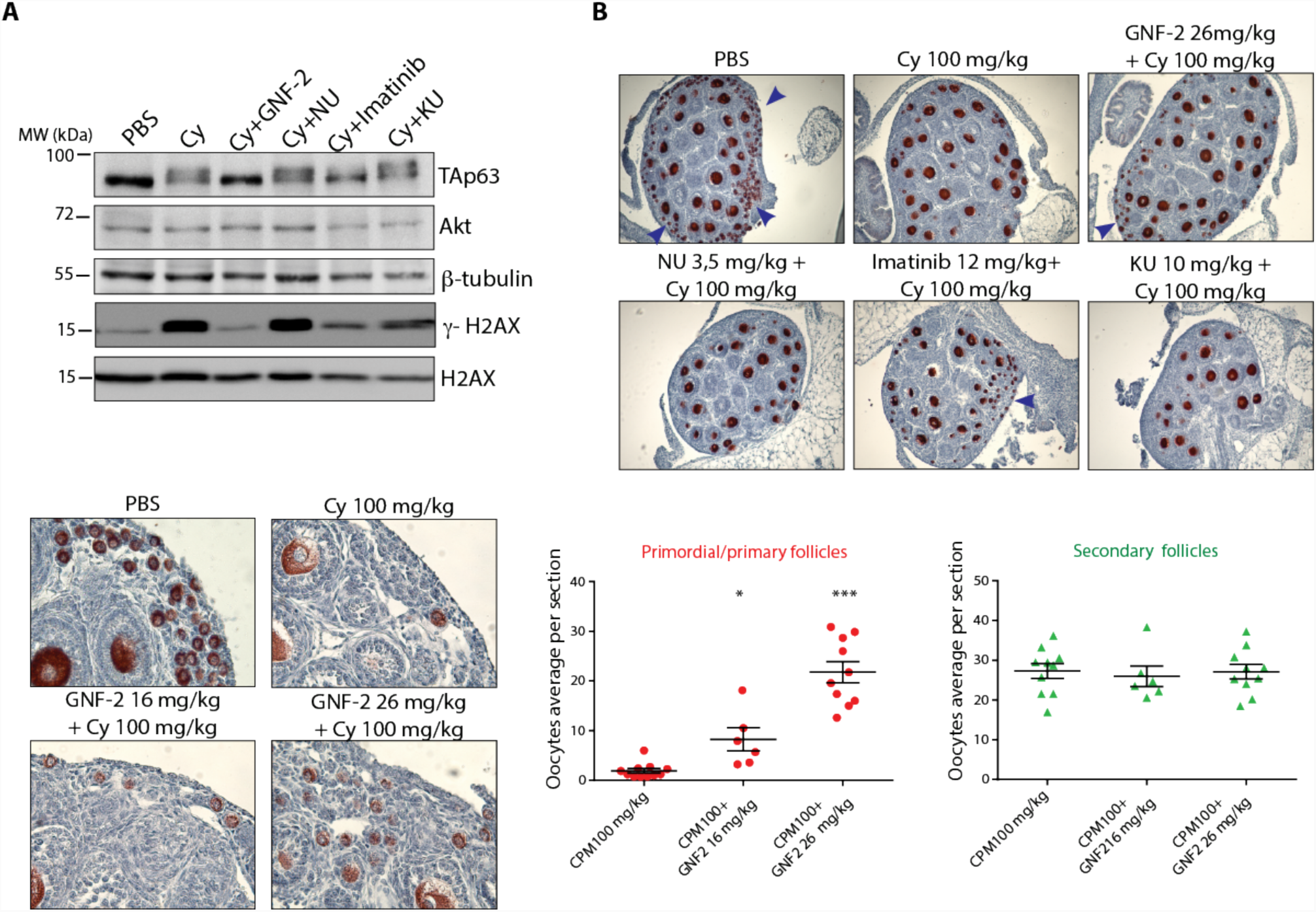
GNF-2 prevents cyclophosphamide-induced loss of reserve follicles. P7 mice were injected with vehicle (PBS) or cyclophosphamide (100 mg/kg) alone or in combination with different kinase inhibitors (GNF-2, imatinib, NU7441, or KU55933). Mice were sacrificed within 24 h from injection. (A) Western blot analysis showed that GNF-2 prevented TAp63 activation/shift and γH2AX phosphorylation, whereas NU7441 and KU55933 were less effective in preventing TAp63 activation/shift. (B, C) Ovaries were dissected 3 days after injection and analyzed with IHC assay using Msy-2 antibody. (D) Several ovaries from three independent experiments were analyzed; each dot in the box plot represents the average number (primordial/primary or secondary) of follicles per section of each gonad collected. Statistical significance was determined using one-way analysis of variance (ANOVA) (*P < 0.05; ***P < 0.001 as compared with the group treated with 100 mg/kg cyclophosphamide). Blue arrows indicate follicle reserves.

### GNF-2 prevents the cyclophosphamide-induced loss of follicle reserve

We dissected and analyzed the cyclophosphamide-treated ovaries by western blotting. Ovarian lysates from mice injected with PBS or cyclophosphamide (100 mg/kg) alone or in combination with kinase inhibitors targeting ABL (GNF-2, imatinib), DNA-PK (NU7441), or ATM (KU55933) revealed the GNF-2-mediated prevention of TAp63 activation (i.e., absence of mobility shift) and γH2AX generation. NU7441 and KU55933 did not prevent TAp63 activation or γH2AX generation (Fig. 5A). On the other hand, imatinib partially prevented TAp63 activation/shift and γH2AX expression. These results were also confirmed by IHC assay performed on the ovaries dissected 3 days after injection (Fig. 5B). Ovarian sections showed a massive depletion in primordial and primary follicles after cyclophosphamide treatment, whereas a significant rescue in follicle reserve was observed in the ovaries co-treated with cyclophosphamide and GNF-2 (Fig. 5C). We counted primordial, primary, and secondary follicles from middle-ovary sections (Fig. 5D), as previously described ^22^. We failed to observe any significant difference in secondary follicles in cyclophosphamide-treated and control groups.

### Imatinib and NU7441 partially mitigate the toxic effect of cyclophosphamide

We evaluated the effect of different doses of two kinase inhibitors imatinib and NU7441. Both compounds bind to the ATP-binding cleft of kinases and are quite selective for their main target kinases. However, imatinib binds to other kinases such as platelet-derived growth factor receptor (PDGFR) and c-KIT^24^, while NU7441 also inhibits phosphatidylinositol-4,5-bisphosphate 3-kinase (PI3K) and has little activity against ATM and ATR^25^. We assessed the effect of these compounds in combination with cyclophosphamide. We injected mice with cyclophosphamide (100 mg/kg) in the presence of each inhibitor. TUNEL assay result showed that the co-treatment with imatinib or NU7441 had milder effects on the prevention of granulosa cell death than co-treatment with GNF-2 (Fig. 6A and Supplementary Fig. 6A). We found that imatinib and NU7441 failed to prevent DNA-PK-γH2AX-TAp63 activation (Fig. 6B and Supplementary Fig. 6B). However, at low dose, imatinib temporarily prevented apoptosis in follicle reserve (Fig. 6C). This observation was also confirmed by IHC assay (Fig. 5B) performed in ovarian sections dissected after 3 days of injection (Fig. 6D). NU7441 and cyclophosphamide co-treatment mildly mitigated apoptosis in follicle reserve (Supplementary Fig. 6C) as confirmed by IHC staining (Fig. 5B) of the ovarian sections dissected 3 days after injection (Supplementary Fig. 6D). In conclusion, imatinib and NU7441 are less effective than the allosteric inhibitor GNF-2 in the prevention of cyclophosphamide-induced follicle loss. This observation may depend on the inhibition of kinases, including the receptor tyrosine kinase c-KIT and PIK3 that play important roles in follicle reserve. On the contrary, no known target for the allosteric compound GNF-2, other than ABL kinases, has been reported so far^26^.

**Figure 6.**
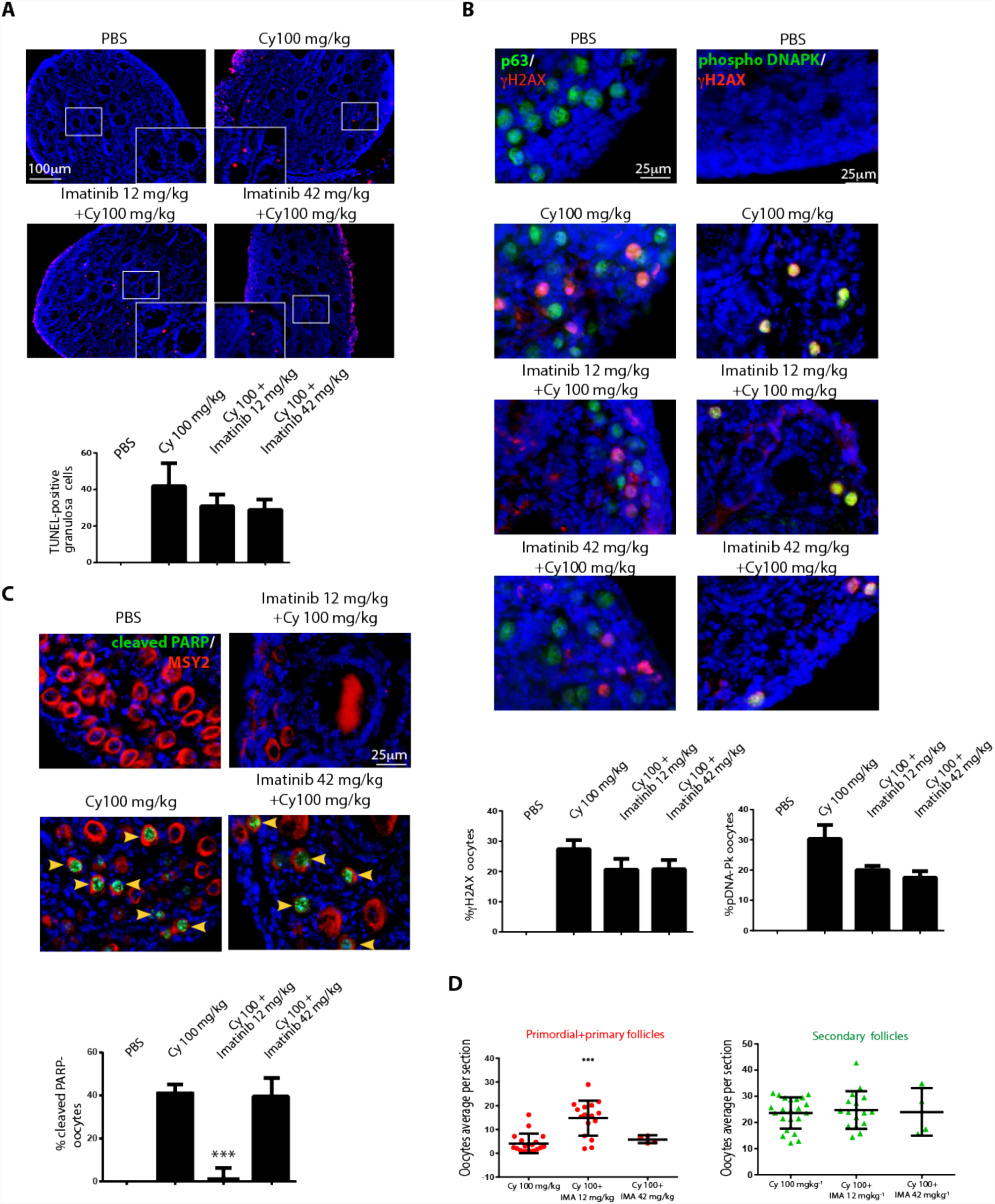
Imatinib partially prevents oocyte apoptosis induced by cyclophosphamide. P7 mice were injected with vehicle (PBS) or cyclophosphamide (100 mg/kg) with/without increasing concentrations of imatinib (12 and 42 mg/kg) and sacrificed within 16 h from injection. (A) Ovarian sections were analyzed by in situ TdT-mediated dUTP nick-end labelling (TUNEL) assay. The graph shows the quantification of TUNEL-positive cells. Quantification of TUNEL-positive cells was performed by counting six different middle ovarian sections derived from three distinct ovaries. (B) γH2AX and DNA-PK activation was observed with IF assay using phospho-specific antibodies and p63 was used as a nuclear marker for germ cells. Quantification was performed by counting several (6 < x < 8) middle ovarian sections derived from three distinct ovaries. Co-staining for pDNA-PK and γH2AX showed the activation of DNA damage response in reserve oocytes. Quantification was performed by counting several (6 < x < 8) middle ovarian sections derived from three distinct ovaries. (C) Ovarian reserve apoptosis was assessed by IF assay using antibodies against cleaved PARP (green) and Msy2 (red), a cytoplasmic antigen of germ cells. Quantification of cleaved PARP-positive cells was performed by counting several (6 < x < 8) middle ovarian sections derived from three distinct ovaries. (A; B; C) Bar column represents mean ± s.d.; statistical significance was determined using one-way analysis of variance (ANOVA) (***P < 0.001 as compared with the group treated with 100 mg/kg cyclophosphamide). (D) Ovaries dissected 3 days after injection were analyzed with IHC assay using Msy2 antibody (Fig. 5B). Ovaries from three independent experiments were analyzed; each dot in the box plot represents the average number of follicles (primordial + primary and secondary) per section of each gonad collected. Statistical significance was determined using one-way analysis of variance (ANOVA) (***P < 0.001 as compared with cyclophosphamide-treated group at 100 mg/kg). Scale bar magnification, 100 μm for TUNEL assay and 25 μm for IF assay.

### GNF-2 prolongs fertility in female mice injected with cyclophosphamide

To investigate the long-term effect of concomitant GNF-2 administration, co-treated mice were allowed to grow and eventually mated with fertile males. Mice were injected at P7, 3 days after few ovaries from each experimental group were analyzed with IHC assay (Fig. 7A and Supplementary Fig. 7A). We also monitored hair recovery for each experimental group after 2 weeks of injection (Supplementary Fig. 7B) and evaluated pubertal ovaries of cyclophosphamide-treated mice by IHC staining before fertility test (Supplementary Fig. 8). In addition, we measured the average weight of mice from each experimental group over 4 weeks from injection (Fig. 7B). We followed the mating capability and the number of pups delivered during six breeding rounds. No evident differences in behavior and development were observed between the off-springs from each experimental group during the first post-natal week. However, we observed that fertility was impaired in the females injected with cyclophosphamide and DPH (Fig. 7C and D). The mice treated with cyclophosphamide and DPH were infertile after three mating rounds, quite before those treated with cyclophosphamide. On the other hand, co-treatment of mice with GNF-2 and cyclophosphamide prolonged the fertility as compared to cyclophosphamide-treated mice and increased the cumulative number of pups compared to cyclophosphamide + DPH and cyclophosphamide groups (Fig. 7E). The gross morphology of the ovaries dissected from adult mice showed a clear difference in size between gonads of each experimental group. The analysis of ovarian sections by IHC assay with Msy2 confirmed the nearly complete absence of follicle reserve in the ovaries from cyclophosphamide + DPH and cyclophosphamide groups (Fig. 7F).

**Figure 7.**
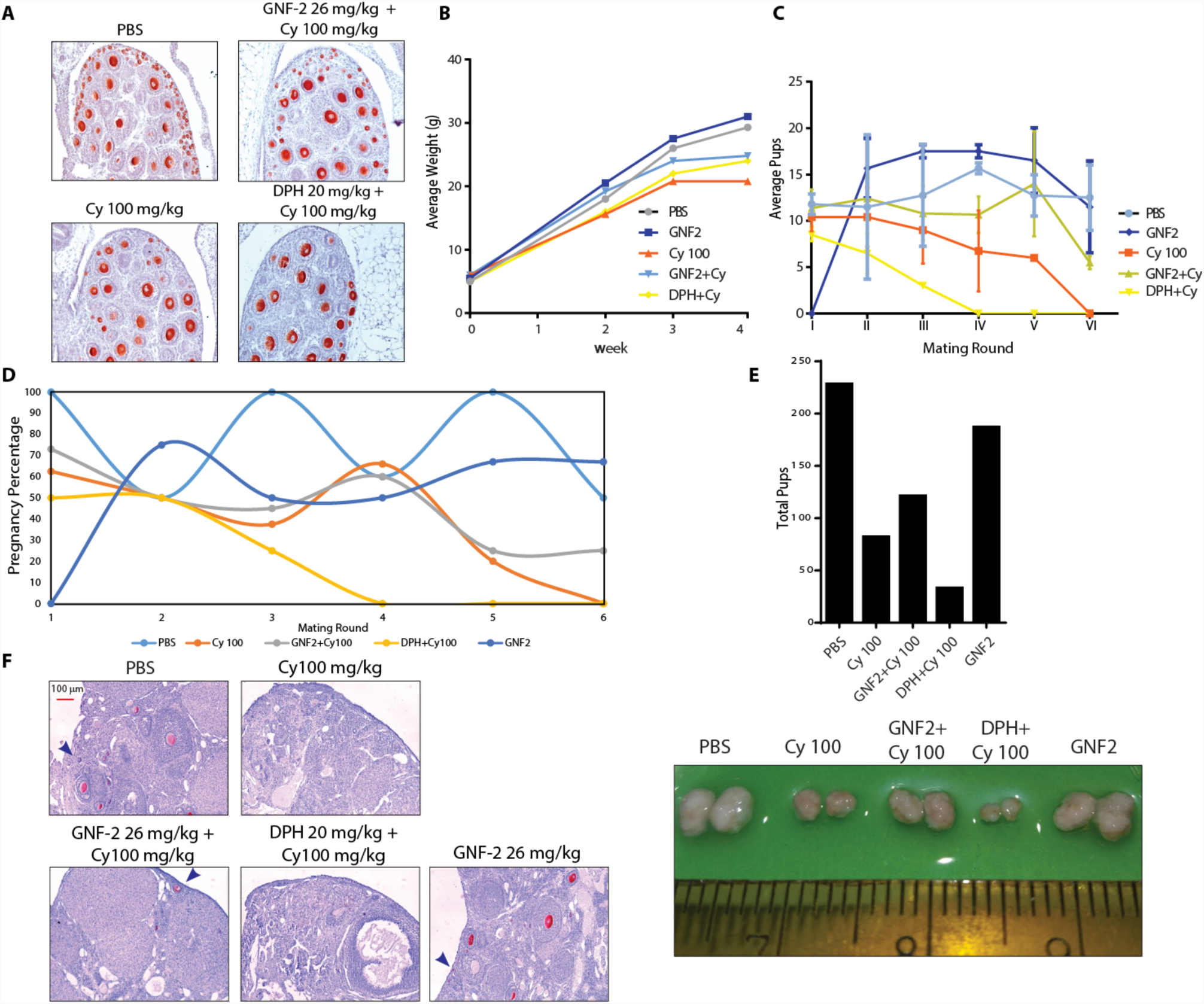
Co-treatment with allosteric compounds modulates fertility in cyclophosphamide-injected mice. (A) Ovaries of each experimental group were dissected 3 days after injection and analyzed with IHC assay using Msy-2 antibody. (B) Graph of average weight following 4 weeks from injection for each experimental group. (C) Time course change in litter size of each group. (D) Percentage of pregnancy rate for the experimental groups throughout mating rounds. (E) Total number of pups delivered by females of each experimental group. (F) Gross morphologies and representative sections of the same ovaries analyzed by IHC assay with Msy2 antibody. Scale bar, 100 μm.

## Discussion

Cyclophosphamide is currently used for the treatment of pediatric cancers. A major concern is the ovarian reserve depletion induced by cyclophosphamide treatment. In the present study, we used a pre-pubertal mouse model to show *in vivo* the signaling axes induced by cyclophosphamide. We also tested small molecules that could limit the cyclophosphamide-induced toxicity by acting as putative ‘ferto-adjuvants.”

Genetically modified mouse models have facilitated the study of the signaling pathways induced in ovary by IR or chemotherapy. Gene targeting approaches may oversimplify the interpretation of the results. The lack of a single node/protein could affect signaling circuits at different levels and the observations reported herein in knockout mice may be attributed to the multiple effects that are difficult to dissect. Small molecules offer unique advantages in terms of interpretation of results because of their transient inhibitory effects on targeted proteins (and/or pathways). Pharmaceutical inhibitors may have a direct relevance with patient care.

Our study showed that cyclophosphamide induced apoptosis in the ovarian reserve through signaling axes involving DNA-PK/(ATM), CHK2, p53, and TAp63*α*. In addition, the co-treatment of cyclophosphamide and GNF-2 affected the DDR induced by cyclophosphamide in the ovary. We also showed that the reserve oocytes rescued from immediate degeneration were healthy enough to produce normal off-springs. Gross morphology and IHC analyses of ovary sections performed either before fertility test or after infertility detection in the first group of treated mice confirmed that the concomitant administration of GNF-2 and cyclophosphamide had long-term effects. Our data are reinforced through the use of an allosteric activator (DPH) that exerted opposite effects as compared to GNF-2. Co-treatment with DPH and cyclophosphamide enhanced the DDR induced by cyclophosphamide in ovaries and shortened mouse fertility. We also compared the effect of the transient administration of inhibitors targeting DDR apical kinases and observed mild protective effects exclusively at low doses of kinase inhibitors. Why allosteric compounds targeting ABL are more effective than ATP-competitive kinase inhibitors in rewiring apoptotic pathways induce by cyclophosphamide is questionable. We hypothesized that either i) the theory of “burnout effect”^13^ or ii) the direct DNA damage of follicle reserve in the ovary assaulted by cyclophosphamide may be associated with this effect. Both theories rely on the fact that the damaged somatic cells communicate with the oocyte. In this scenario, allosteric ABL compounds could affect follicle apoptosis by targeting the transmission of stress signals from pre-granulosa cells to the oocyte. Although the molecular details remain unclear, our current model suggests that ABL kinases may act as “allosteric devices” that contribute to fuel the molecular events underlying stress signaling/DDR in the follicles^27^. This model is supported by the observation that GNF-2 prevents the activation of the AKT-FOXO3 signaling axis induced by cyclophosphamide. We also found that DPH promotes the activation of the AKT-FOXO3 signaling axis following cyclophosphamide treatment and that the reserve oocytes were positive for both p-AKT/p-FOXO3 and γH2AX expression. Thus, the presence of the early marker of DDR is consistent with the activation of the AKT-FOXO3 pathway. These observations suggest that oocytes leave their follicular quiescence to initiate the DNA damage signaling pathways that end in oocyte apoptosis. Whether this is a general mechanism occurring with other DNA-damaging chemotherapeutic drugs and IR is yet unknown.

Evidences suggest that ABL kinase inhibitors protect the ovarian reserve from cisplatin-induced degeneration^19, 20, 21, 22^. However, the mechanism underlying the imatinib-mediated protection of oocytes remains debated. Genetic studies have shown that the oocyte ABL kinases are dispensable for cisplatin-induced TAp63*α* activation^10^. However, ABL kinases in complex with inhibitors may not recapitulate the results observed in ABL null mice, given that ABL proteins take part to opposing signaling pathways in cells^28^.

Recent studies have identified CHK2 and p53 as two important players in the efficient removal of oocytes with unrepaired meiotic DNA DBSs^8^. In addition, CHK2-dependent TAp63*α* phosphorylation is a key event induced in response to IR^6, 8^. We show that cyclophosphamide induced DNA damage checkpoint pathways that involve signaling of DNA-PK to CHK2, which in turn communicates with p53 and TAp63, revising a previously proposed model^8^. We found p-p53 (Ser15) in the nucleus of the damaged oocytes, supporting the direct role of p53 in driving the apoptotic response of follicles. This observation is intriguing, as TAp63*α* is considered as a key player in quality control of germ cells. However, activated TAp63*α* exerts modest activity as a transcription factor as compared with p53; thus, the quality control and apoptotic response induced by cyclophosphamide require the concomitant outcomes of both factors.

A recent work showed that mouse models lacking PUMA (a pro-apoptotic protein) retained fertility after chemotherapy^29^. This observation supports the results that reserve oocytes from PUMA^-/-^ mice may sufficiently repair themselves to support healthy off-springs. These findings strengthen the argument that oocytes are in fact capable of efficient DNA repair in response to the inhibition of the apoptotic pathway in the ovary^30^.

The concomitant administration of GNF-2 and cyclophosphamide results in the inhibition of the AKT-FOXO3 signaling axis and DDR, thereby preventing cyclophosphamide-induced apoptosis of ovarian reserve. Thus, the feasibility of ferto-adjuvant therapies based on allosteric ABL compounds is established. With this in mind, we should consider that the systemic administration of allosteric inhibitors may interfere with the efficacy of cancer therapies. Clinical applications may warrant studies to improve the targeted delivery of these compounds to the gonads. Despite these efforts, the discovery of Asciminib (ABL001)^31, 32^, the first allosteric ABL inhibitor used in clinic, offers promises to develop ferto-protective strategies suitable for pediatric cancer patients.

## Acknowledgments

We thank Gianni Cesareni and Luisa Castagnoli for their valuable support. We are indebted to Stefano Cannata, Stefano Pirrò, and Sergio Bernardini for their technical advice. We are grateful to Gianni Cesareni and Luca Jovine for their helpful comments on the manuscript. This work was supported by grants from AIRC (IG11344) to S.G.

## Author Contributions

G.B. performed the majority of the experiments with the help of V.I. S.C. performed confocal analysis and contributed to follicle quantification. L.M. performed IF assays for p-CHK2 and p-53 as well as the fertility test. E.M. performed all experiments shown in Supplementary Fig. 4. V.V. performed IHC assay with adult ovaries and follicle quantification shown in Supplementary Fig. 8. M.D. provided support and reagents and helped with the critical reading of the manuscript. S.G. designed and directed the study, wrote the manuscript.

---

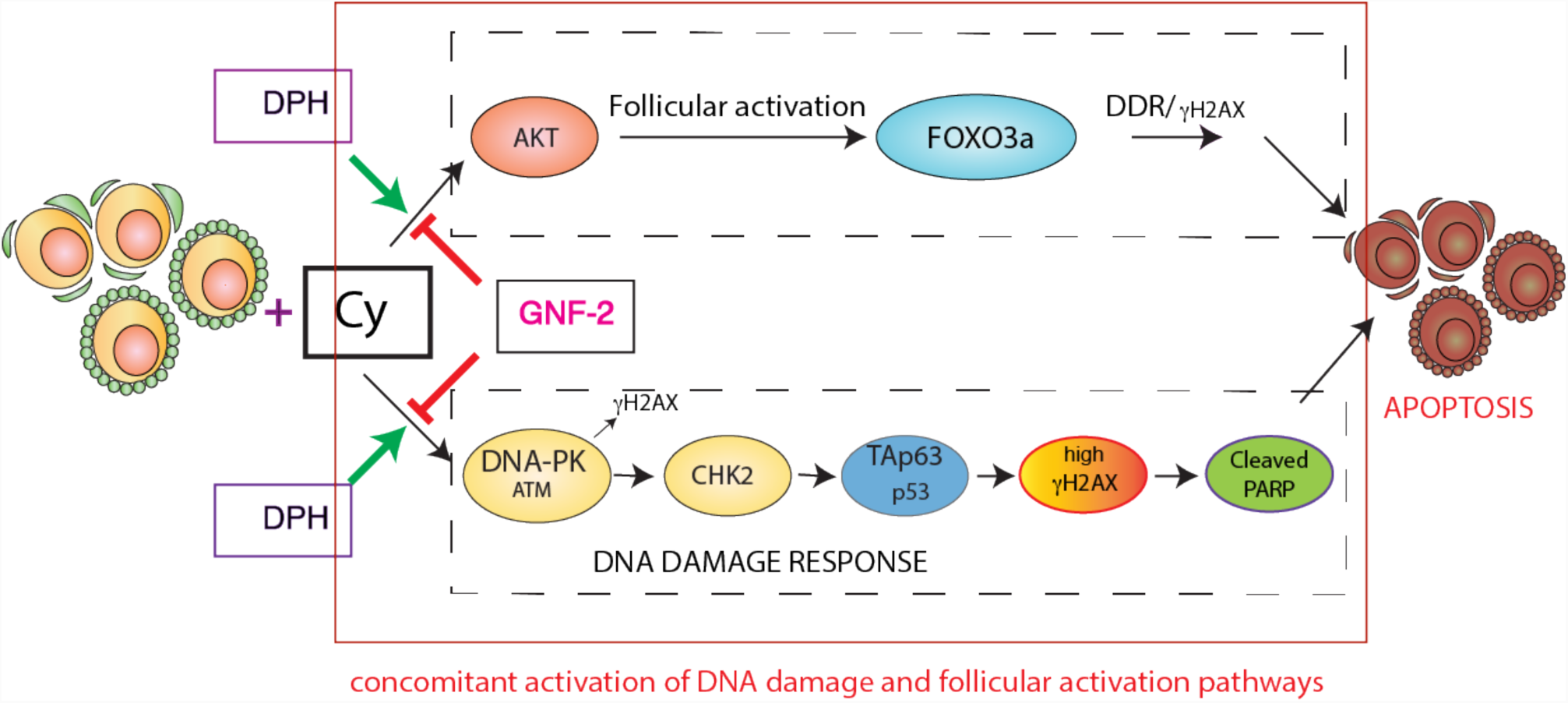
Model: Concomitant administration of GNF-2 and cyclophosphamide results in the inhibition of the AKT-FOXO3 signaling axis and DDR, thereby preventing cyclophosphamide-induced apoptosis of oocyte reserve. On the contrary, DPH enhanced the activation of the AKT-FOXO3 signaling axis and DDR response following cyclophosphamide injection. Oocyte reserve was positive for both p-AKT/p-FOXO3 and γH2AX; thus, the presence of the early marker of DDR was consistent with the activation of the AKT-FOXO3 pathway. This observation suggests that oocytes leave their follicular quiescence to initiate the DNA damage signaling pathway that leads to oocyte apoptosis.

## Notes

**Conflict of Interest statement**: the authors declare no competing financial interests.

## References

1. Morgan S, Anderson RA, Gourley C, Wallace WH, Spears N. How do chemotherapeutic agents damage the ovary? Hum Reprod Update 2012, 18(5): 525–535.

2. Oktem O, Oktay K. Quantitative assessment of the impact of chemotherapy on ovarian follicle reserve and stromal function. Cancer 2007, 110(10): 2222–2229.

3. Meirow D, Biederman H, Anderson RA, Wallace WH. Toxicity of chemotherapy and radiation on female reproduction. Clin Obstet Gynecol 2010, 53(4): 727–739.

4. Suh EK, Yang A, Kettenbach A, Bamberger C, Michaelis AH, Zhu Z, et al. p63 protects the female germ line during meiotic arrest. Nature 2006, 444(7119): 624–628.

5. Livera G, Petre-Lazar B, Guerquin MJ, Trautmann E, Coffigny H, Habert R. p63 null mutation protects mouse oocytes from radio-induced apoptosis. Reproduction 2008, 135(1): 3–12.

6. Deutsch GB, Zielonka EM, Coutandin D, Weber TA, Schafer B, Hannewald J, et al. DNA damage in oocytes induces a switch of the quality control factor TAp63alpha from dimer to tetramer. Cell 2011, 144(4): 566–576.

7. Kerr JB, Hutt KJ, Michalak EM, Cook M, Vandenberg CJ, Liew SH, et al. DNA damage-induced primordial follicle oocyte apoptosis and loss of fertility require TAp63-mediated induction of Puma and Noxa. Molecular cell 2012, 48(3): 343–352.

8. Bolcun-Filas E, Rinaldi VD, White ME, Schimenti JC. Reversal of female infertility by Chk2 ablation reveals the oocyte DNA damage checkpoint pathway. Science 2014, 343(6170): 533–536.

9. Rinaldi VD, Hsieh K, Munroe R, Bolcun-Filas E, Schimenti JC. Pharmacological Inhibition of the DNA Damage Checkpoint Prevents Radiation-Induced Oocyte Death. Genetics 2017, 206(4): 1823–1828.

10. Kim SY, Nair DM, Romero M, Serna VA, Koleske AJ, Woodruff TK, et al. Transient inhibition of p53 homologs protects ovarian function from two distinct apoptotic pathways triggered by anticancer therapies. Cell death and differentiation 2018.

11. Tuppi M, Kehrloesser S, Coutandin DW, Rossi V, Luh LM, Strubel A, et al. Oocyte DNA damage quality control requires consecutive interplay of CHK2 and CK1 to activate p63. Nat Struct Mol Biol 2018, 25(3): 261–269.

12. Bedoschi G, Navarro PA, Oktay K. Chemotherapy-induced damage to ovary: mechanisms and clinical impact. Future Oncol 2016, 12(20): 2333–2344.

13. Kalich-Philosoph L, Roness H, Carmely A, Fishel-Bartal M, Ligumsky H, Paglin S, et al. Cyclophosphamide Triggers Follicle Activation and “Burnout”; AS101 Prevents Follicle Loss and Preserves Fertility. Sci Transl Med 2013, 5(185): 185ra162.

14. Jayasinghe YL, Wallace WHB, Anderson RA. Ovarian function, fertility and reproductive lifespan in cancer patients. Expert Rev Endocrinol Metab 2018, 13(3): 125–136.

15. Kim SY, Kim SK, Lee JR, Woodruff TK. Toward precision medicine for preserving fertility in cancer patients: existing and emerging fertility preservation options for women. J Gynecol Oncol 2016, 27(2): e22.

16. Roness H, Kashi O, Meirow D. Prevention of chemotherapy-induced ovarian damage. Fertil Steril 2016, 105(1): 20–29.

17. Woodruff TK. Preserving fertility during cancer treatment. Nat Med 2009, 15(10): 1124–1125.

18. Shah JS, Sabouni R, Cayton Vaught KC, Owen CM, Albertini DF, Segars JH. Biomechanics and mechanical signaling in the ovary: a systematic review. J Assist Reprod Genet 2018, 35(7): 1135–1148.

19. Gonfloni S, Di Tella L, Caldarola S, Cannata SM, Klinger FG, Di Bartolomeo C, et al. Inhibition of the c-Abl-TAp63 pathway protects mouse oocytes from chemotherapy-induced death. Nature medicine 2009, 15(10): 1179–1185.

20. Kim SY, Cordeiro MH, Serna VA, Ebbert K, Butler LM, Sinha S, et al. Rescue of platinum-damaged oocytes from programmed cell death through inactivation of the p53 family signaling network. Cell death and differentiation 2013.

21. Morgan S, Lopes F, Gourley C, Anderson RA, Spears N. Cisplatin and doxorubicin induce distinct mechanisms of ovarian follicle loss; imatinib provides selective protection only against cisplatin. PloS one 2013, 8(7): e70117.

22. Maiani E, Di Bartolomeo C, Klinger FG, Cannata SM, Bernardini S, Chateauvieux S, et al. Reply to: Cisplatin-induced primordial follicle oocyte killing and loss of fertility are not prevented by imatinib. Nat Med 2012, 18(8): 1172–1174.

23. Yang J, Campobasso N, Biju MP, Fisher K, Pan XQ, Cottom J, et al. Discovery and characterization of a cell-permeable, small-molecule c-Abl kinase activator that binds to the myristoyl binding site. Chemistry & biology 2011, 18(2): 177–186.

24. Buchdunger E, Cioffi CL, Law N, Stover D, Ohno-Jones S, Druker BJ, et al. Abl protein-tyrosine kinase inhibitor STI571 inhibits in vitro signal transduction mediated by c-kit and platelet-derived growth factor receptors. J Pharmacol Exp Ther 2000, 295(1): 139–145.

25. Tavecchio M, Munck JM, Cano C, Newell DR, Curtin NJ. Further characterisation of the cellular activity of the DNA-PK inhibitor, NU7441, reveals potential cross-talk with homologous recombination. Cancer Chemother Pharmacol 2012, 69(1): 155–164.

26. Adrian FJ, Ding Q, Sim T, Velentza A, Sloan C, Liu Y, et al. Allosteric inhibitors of Bcr-abl-dependent cell proliferation. Nature chemical biology 2006, 2(2): 95–102.

27. Gonfloni S. Defying c-Abl signaling circuits through small allosteric compounds. Front Genet 4. 2014, 5: 392.

28. Maiani E, Diederich M, Gonfloni S. DNA damage response: the emerging role of c-Abl as a regulatory switch? Biochemical pharmacology 2011, 82(10): 1269–1276.

29. Nguyen QN, Zerafa N, Liew SH, Morgan FH, Strasser A, Scott CL, et al. Loss of PUMA protects the ovarian reserve during DNA-damaging chemotherapy and preserves fertility. Cell Death Dis 2018, 9(6): 618.

30. Stringer JM, Winship A, Liew SH, Hutt K. The capacity of oocytes for DNA repair. Cell Mol Life Sci 2018.

31. Wylie AA, Schoepfer J, Jahnke W, Cowan-Jacob SW, Loo A, Furet P, et al. The allosteric inhibitor ABL001 enables dual targeting of BCR-ABL1. Nature 2017, 543(7647): 733–737.

32. Schoepfer J, Jahnke W, Berellini G, Buonamici S, Cotesta S, Cowan-Jacob SW, et al. Discovery of asciminib (ABL001), an allosteric inhibitor of the tyrosine kinase activity of BCR-ABL1. J Med Chem 2018.

