## Supplementary for "Kinase-independent inhibition of cyclophosphamide-induced pathways protects the ovarian reserve and prolongs fertility"

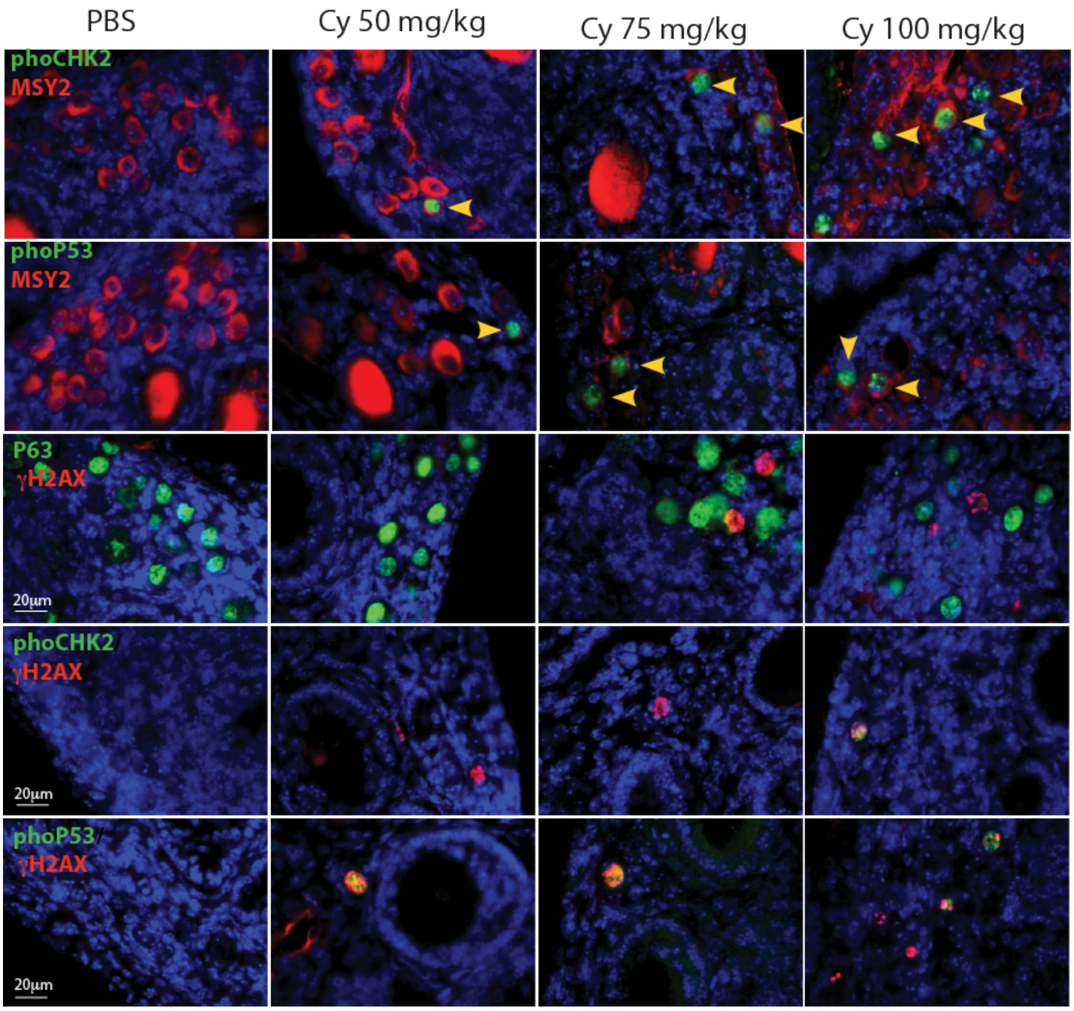


**Supplementary Fig.1 Cyclophosphamide induces Chk2 and p53 activation in nucleus of reserve oocytes.**

P8 mice were injected with vehicle (PBS) or increasing concentrations of cyclophosphamide (50, 75, 100mg/kg) and sacrificed within 16 h from injection. IF assay with two specific phospho-antibodies for Chk2, or p53 (green) or and Msy2 (red), a cytoplasmic antigen of germ cells. Co-staining of p-Chk2 and γH2AX or p53 and γH2AX shows the activation of DNA damage response pathway in reserve oocytes. Yellow arrows at the bottom indicate positive oocytes. Scale bar, magnification 20µm.

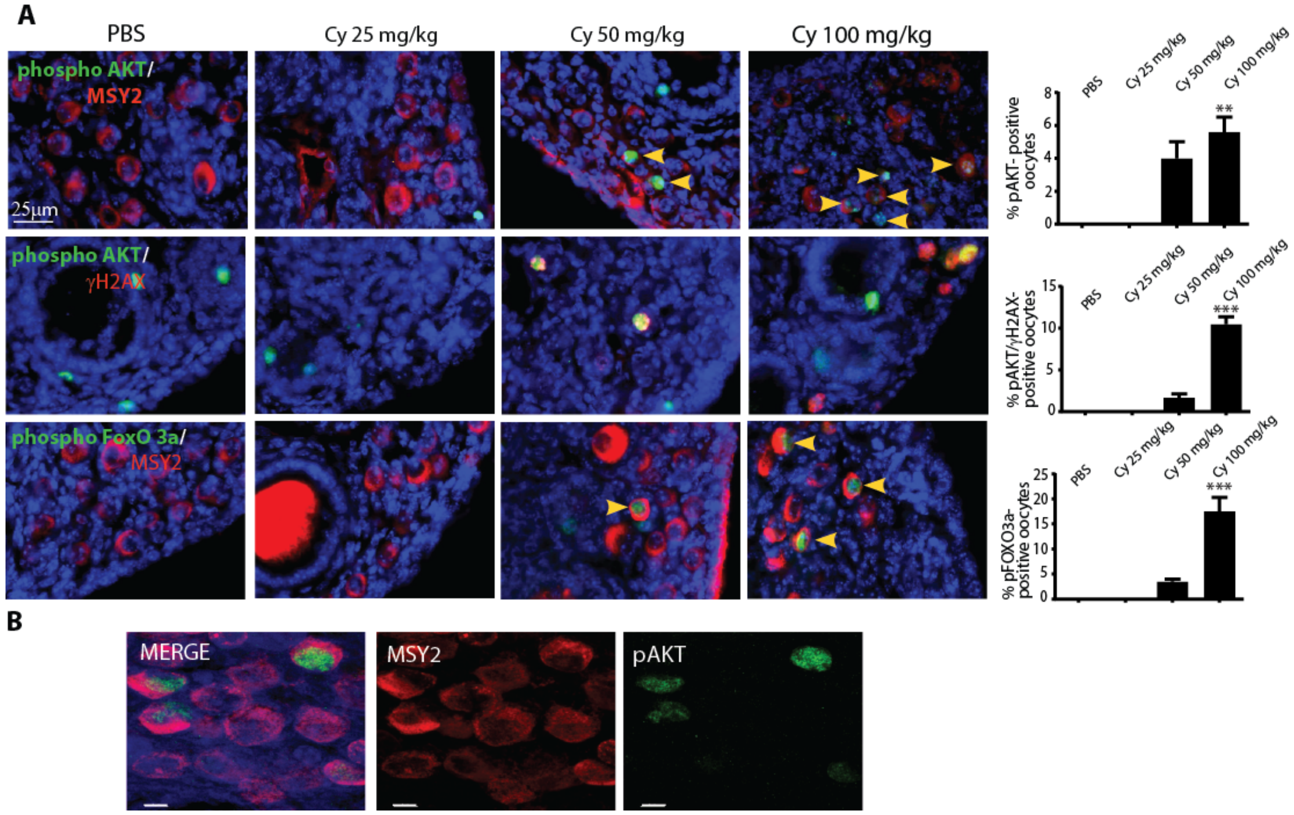


**Supplementary Fig. 2. Cyclophosphamide induces the AKT- FOXO3 pathway in the nucleus of the reserve oocyte.**

P7 mice were injected with vehicle (PBS) or increasing concentrations of cyclophosphamide (25, 50 and 100 mg/kg) and sacrificed within 16 h from injection. AKT(T308) and FOXO3(S253) phosphorylation are followed by IF assay with a phospho-specific antibody (green) and Msy2 (red), a cytoplasmic antigen of germ cells. Yellow arrows indicate oocytes positive for AKT (upper panel) and FOXO3a (lower panel). In the central panel, co-staining of p-AKT and γH2AX shows the activation of DNA damage response pathway in reserve oocytes. Quantification was obtained by counting several (6<x<8) middle ovarian sections derived from distinct ovaries. (B) Confocal images of middle ovarian sections from cyclophosphamide -injected mice confirm the presence of p-AKT in the nucleus of germ cells. Scale bar magnification 25μm for IF assay and 7μm for confocal images. (A) Bar column represents mean ± s.d.; statistical significance was determined using one-way analysis of variance (ANOVA) (**P<0.01; ***P<0.001 as compared with PBS-treated group).

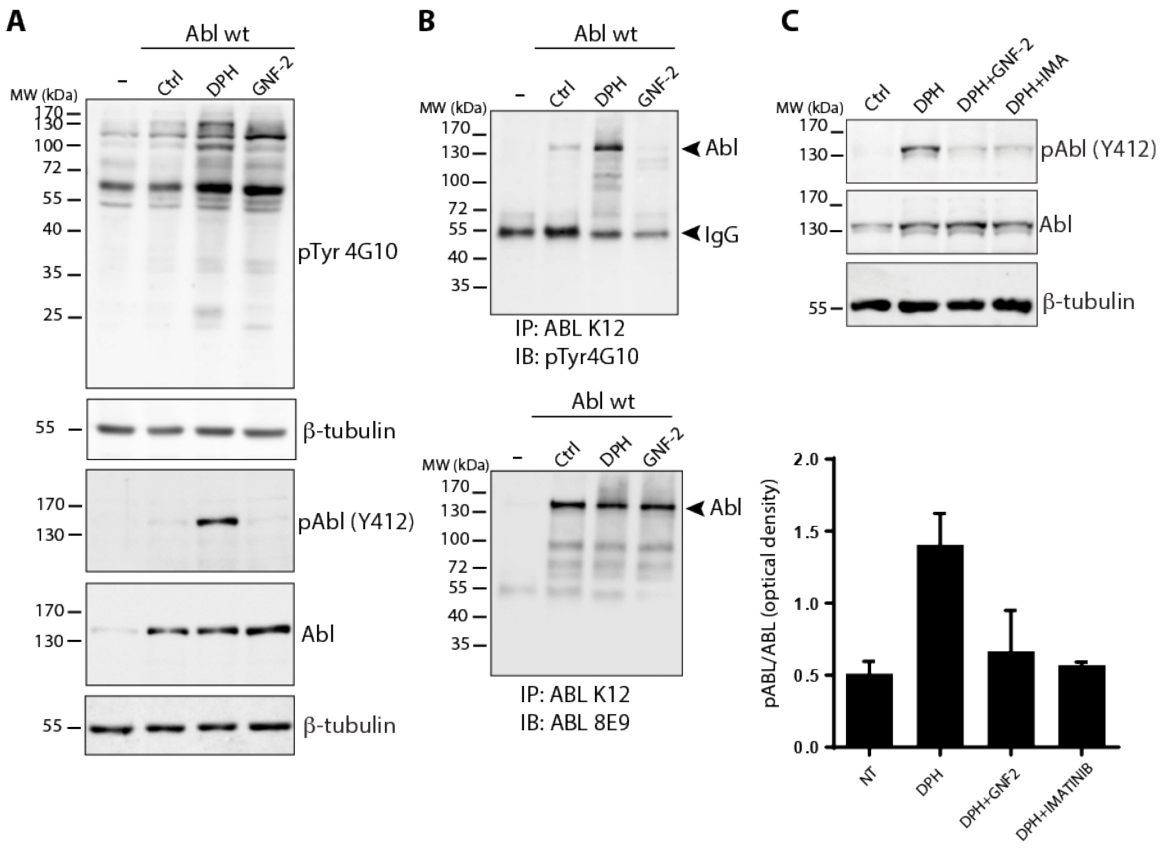


**Supplementary Fig.3 Allosteric compounds modulate ABL catalytic activity in H1299**
Human lung carcinoma cell line H1299 was used for *in vitro* studies to monitor the ABL kinase activity following GNF-2, DPH and Imatinib exposure. (A) Western blotting of total lysate from H1299 cells transfected with a plasmid encoding wild type ABL. DPH treatment induces auto-phosphorylation of Y412 in the activation loop of ABL kinase, while GNF-2 has an opposite effect. (B) Immunoprecipitation assay with ABL polyclonal antibodies was done with the same H1299 extracts to further confirm the data. (C)Total lysates fromH1299 cells, treated with DPH or GNF-2, show that the allosteric compounds affect the kinase activity of endogenous ABL.

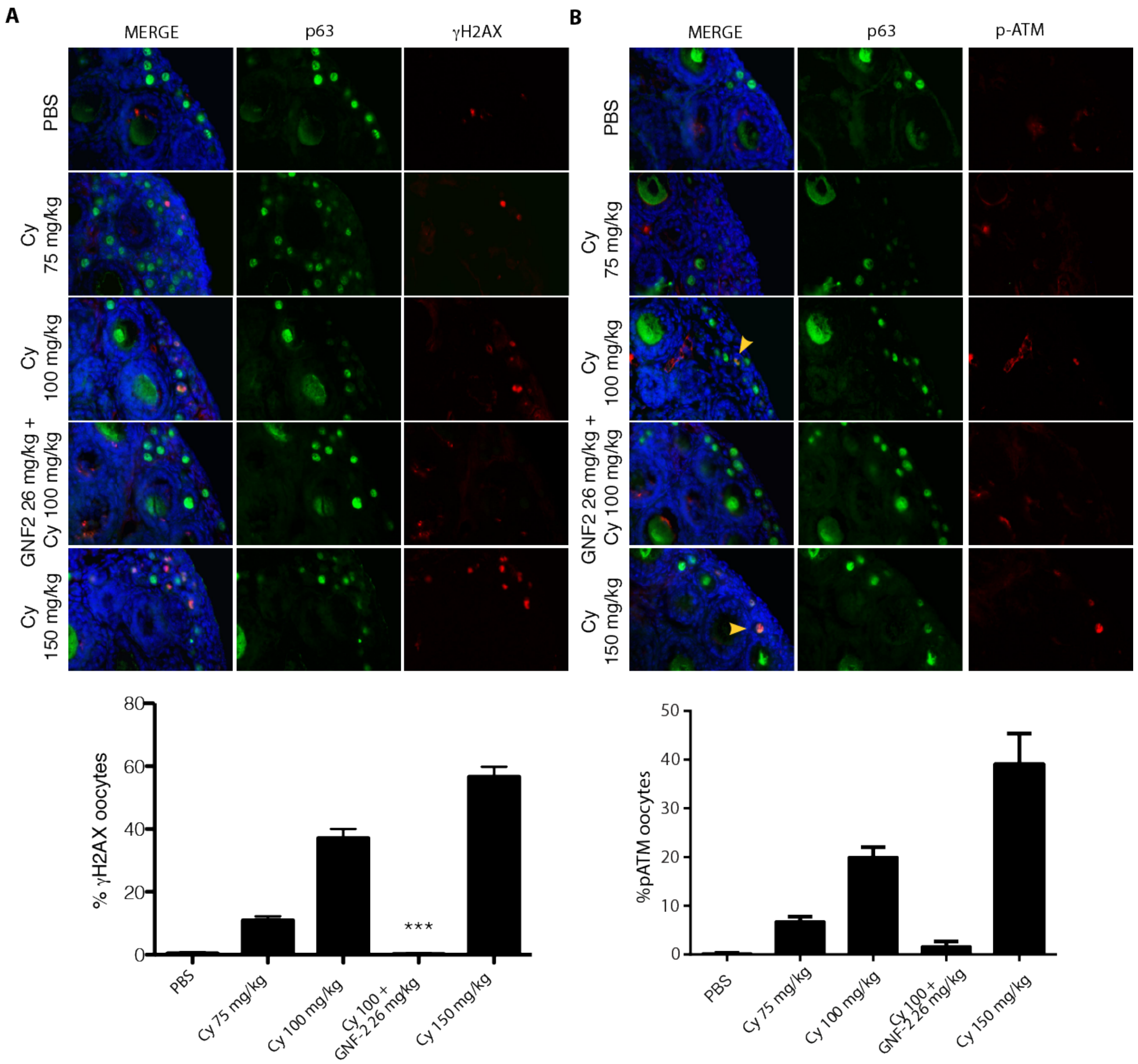


**Supplementary Fig.4 GNF-2 affects ATM phosphorylation in the follicle reserve**

P8 mice were injected with vehicle (PBS) or increasing concentration of cyclophosphamide (75, 100 and 150 mg/kg). Cyclophosphamide was injected alone or in presence of GNF-2. Co-staining of p63 (green) and γH2AX or phosphorylated ATM (red) shows an increase in the number of oocytes positive to γH2AX as indicated in the graph (A) Oocytes with a reduced level of TAp63 show a high expression of phosphorylated ATM. (B) Co-treatment with GNF-2 prevents both ATM activation and γH2AX in the reserve oocytes. Quantification was performed by counting several (6<x<8) middle ovarian sections per condition. Bar column represents mean ± s.d.; statistical significance was determined using one-way analysis of variance (ANOVA) (**P<0.01; ***P<0.001 as compared to 100 mg/kg cyclophosphamide).

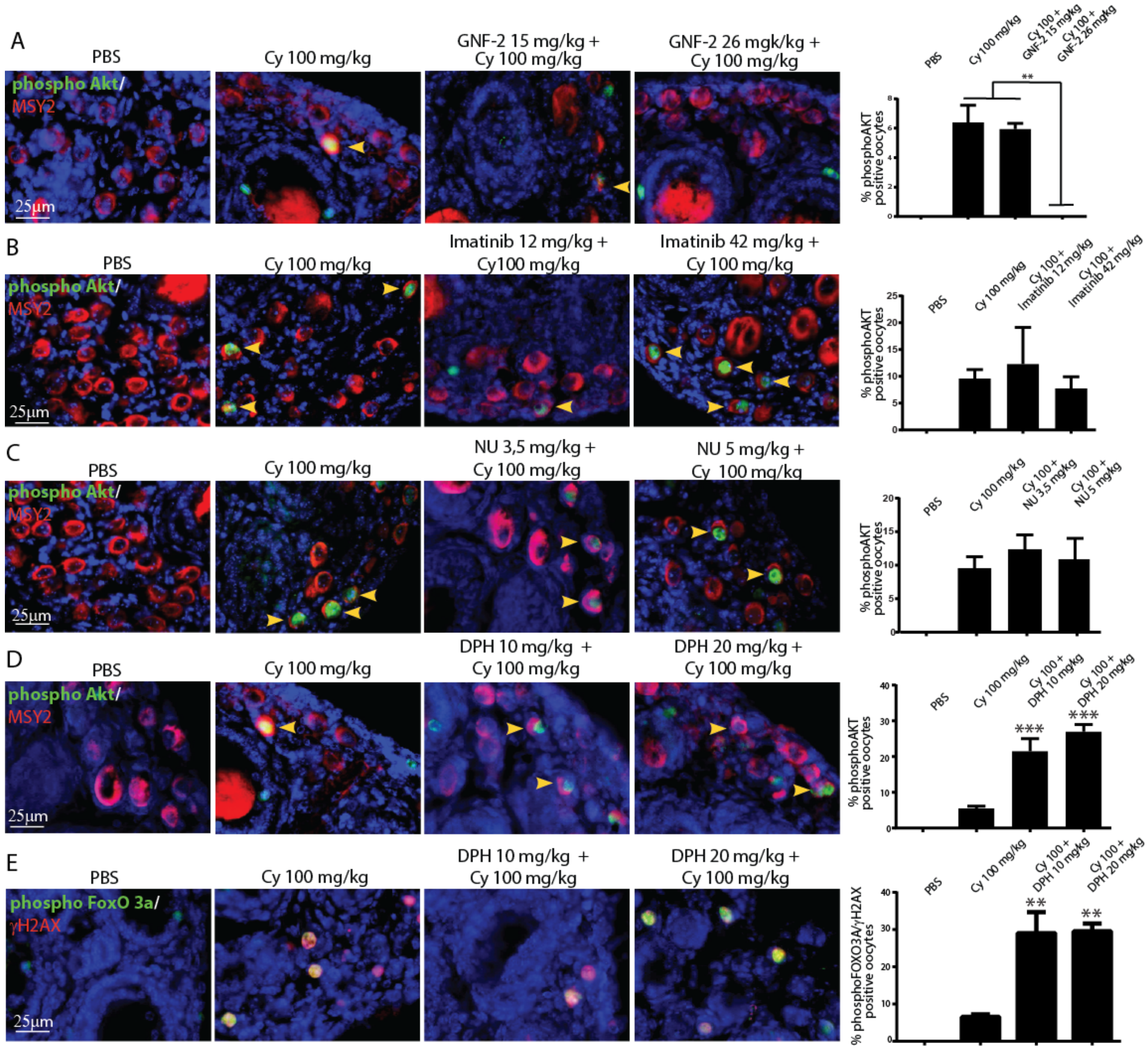


**Supplementary Fig.5 Allosteric compounds modulate AKT-FOXO3 signaling axis induced by cyclophosphamide in the nucleus of reserve oocytes.**

P7 mice were injected with vehicle (PBS) or cyclophosphamide (100 mg/kg) with/out increasing concentration of GNF2 (A), Imatinib (B), NU7441 (C) or DPH (D) and were sacrificed within 16-24 h from injection. Ovarian sections were analyzed by IF assay with specific phospho-antibodies for AKT (T308) (green), FOXO3a(S253) (green), Msy2 (red), or γH2AX (S139). A) GNF2 (26 mg/kg) co-treatment prevents the phosphorylation of AKT in the nucleus of reserve oocytes; B, C) While, Imatinib or NU7441 (DNA-PK inhibitor) do not prevent AKT phosphorylation induced by cyclophosphamide; D) DPH co-treatment enhances the phosphorylation of AKT and well as the phosphorylation of FOXO3a. Quantification was performed obtained by counting several (6<x<10) middle ovarian sections derived from three distinct ovaries. Scale bar magnification 25μm. Bar column represents mean ± s.d., statistical significance was determined using one-way analysis of variance (ANOVA) (**P<0.01; ***P<0.001 as compared to 100mg/kg cyclophosphamide).

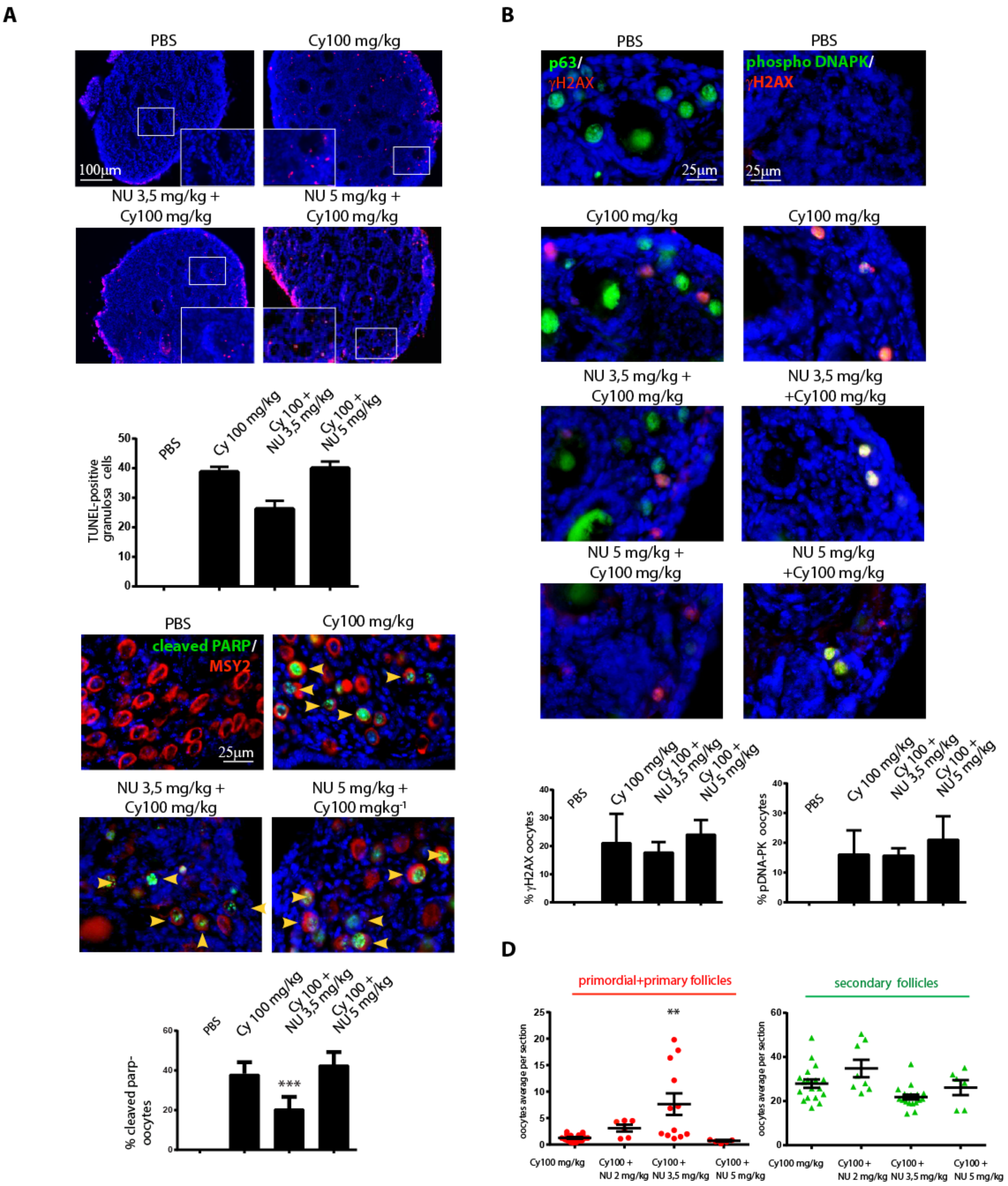
A
PBS
Cy100 mg/kg
100μm
NU 3,5 mg/kg +
Cy100 mg/kg
NU 5 mg/kg +
Cy100 mg/kg
TUNEL-positive granulosa cells
PBS
Cy 100 mg/kg
Cy 100 + NU 3,5 mg/kg
Cy 100 + NU 5 mg/kg
B
PBS
p63/ γH2AX
phospho DNAPK/ γH2AX
25μm
Cy100 mg/kg
NU 3,5 mg/kg + Cy100 mg/kg
NU 5 mg/kg + Cy100 mg/kg
% γH2AX oocytes
% pDNA-PK oocytes
C
PBS
cleaved PARP/ MSY2
Cy100 mg/kg
25μm
NU 3,5 mg/kg + Cy100 mg/kg
NU 5 mg/kg + Cy100 mgkg-1
% cleaved parp-oocytes
***
D
primordial+primary follicles
secondary follicles
oocytes average per section
**
Cy100 mg/kg
Cy100 + NU 2 mg/kg
Cy100 + NU 3,5 mg/kg
Cy100 + NU 5 mg/kg

**Supplementary Fig.6 NU7441 does not prevents oocytes loss induced by cyclophosphamide**

P7 mice were injected with vehicle (PBS) or cyclophosphamide (100 mg/kg) with/out increasing concentration of NU7441 (2 mg/kg, 3,5 mg/kg and 5.3 mg/kg) and were sacrificed within 16-24 h from injection. (A) Ovarian sections were analyzed by in situ TdT-mediated dUTP nick-end labelling (TUNEL) assay. The graph shows the quantification of TUNEL-positive cells. Quantification of TUNEL-positive cells was performed by counting six different middle ovarian sections derived from three distinct ovaries. (B) γH2AX and DNA-PK activation was observed with IF assay using phospho-specific antibodies and p63 was used as a nuclear marker for germ cells. Quantification was performed by counting several (6<x<8) middle ovarian sections derived from three distinct ovaries. Co-staining of pDNA-PK and γH2AX showed the activation of DNA damage response in reserve oocytes. Quantification was performed by counting several (6<x<8) middle ovarian sections derived from three distinct ovaries. (C) Ovarian reserve apoptosis was assessed by IF assay using with two specific antibodies cleaved PARP (green) and Msy2 (red), a cytoplasmic antigen of germ cells. Quantification of cleaved PARP-positive cells was obtained by counting several (6<x<8) middle ovarian sections derived from three distinct ovaries. (A; B; C) Bar column represents mean ± s.d., statistical significance was determined using one-way analysis of variance (ANOVA) (***P<0.001 compared with the group treated with 100 mg/kg cyclophosphamide) (D) Ovaries dissected 3 days after injection were analyzed by IHC assay using Msy2 antibody (see Fig.5B). Ovaries from three independent experiments were analyzed; each dot in the box plot represents the average number of follicles (primordial + primary and secondary) per section of each gonad collected. Statistical significance was determined using one-way analysis of variance (ANOVA) (***P*<0.01 ****P*<0.001 as compared with cyclophosphamide-treated group at 100 mg/kg).

Scale bar magnification, 100μm for TUNEL assay and 25 μm for IF assay.

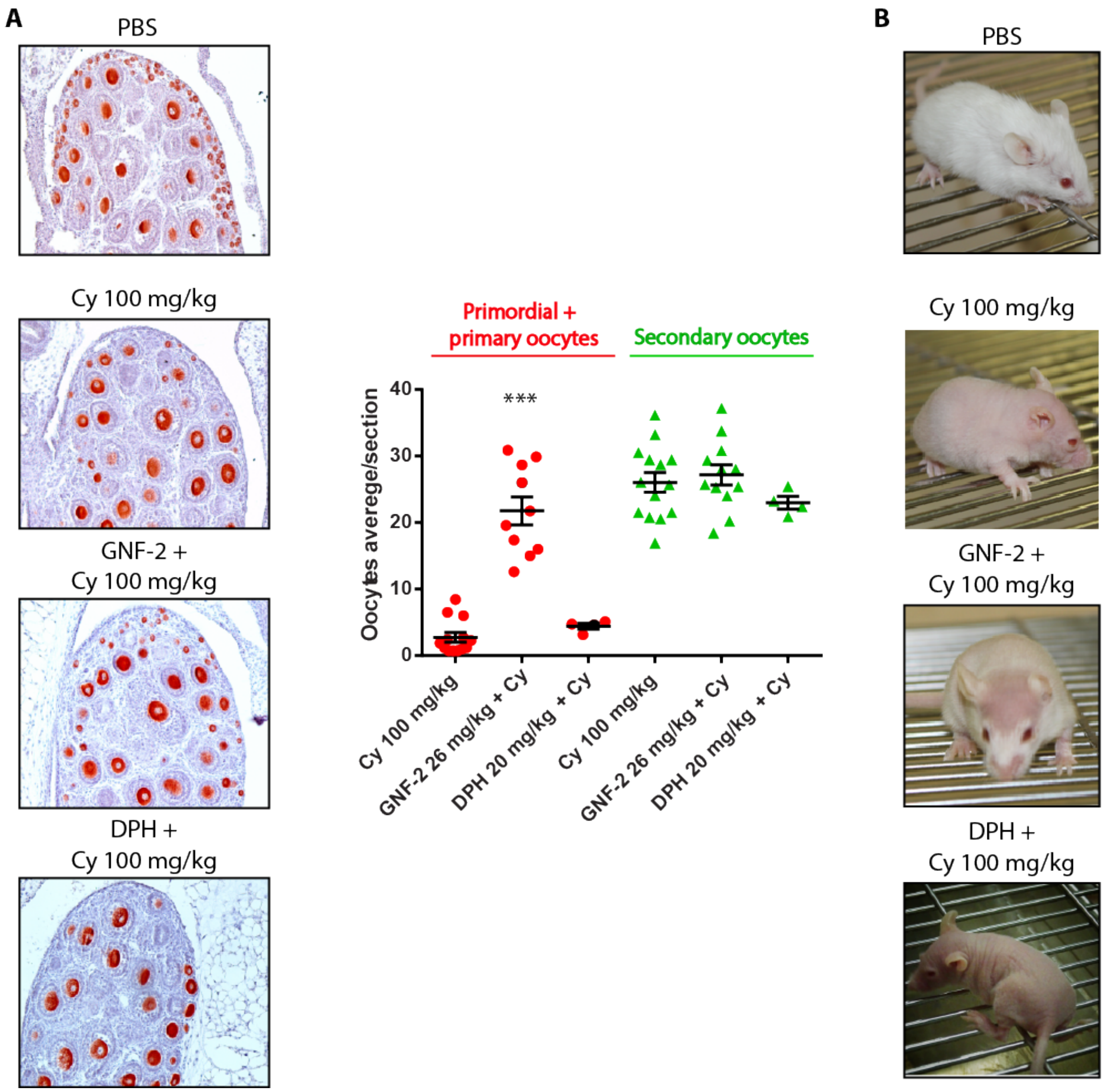


**Supplementary 7. Effects of concomitant administration of allosteric ABL compounds in cyclophosphamide-treated mice**

(A) Ovaries of each experimental group were dissected 3 days after injection and analyzed by IHC assay using Msy-2 antibody. Several ovaries from independent experiments were analyzed, each dot in the box plot represents the average number of follicles (primordial + primary and secondary) per section of each gonad collected. Statistical significance was determined using one-way analysis of variance (ANOVA) (***P<0.001 as compared with cyclophosphamide-treated group at 100 mg/kg). (B) Mice photos were taken about two weeks after injection.

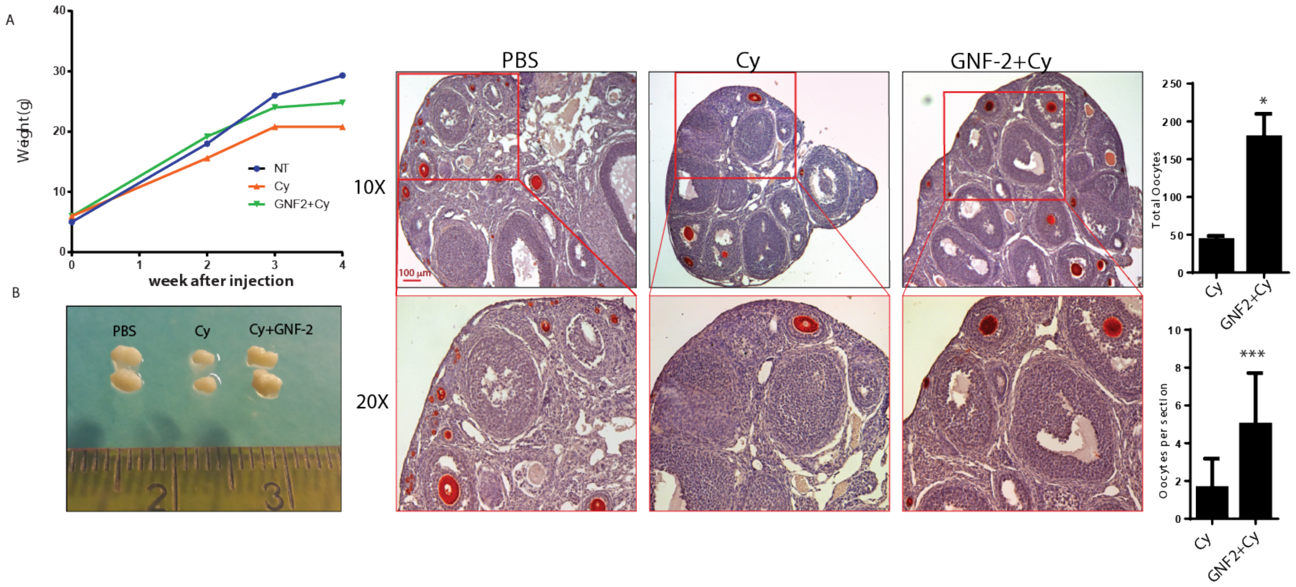


**Supplementary 8. Long-term protection of GNF-2 is evaluated in pubertal ovaries**

(A) Graph of average weight following 4 weeks from injection for each experimental group. (B) Gross morphologies of ovaries dissected from pubertal adult mice before the fertility test. (C) Representative sections of the same ovaries analyzed by IHC assay with Msy2 antibody. Quantification of follicle reserve is also shown. Scale bar, 100µm. Bar column represents mean ± s.d. (n=4). Statistical significance was determined by unpaired Student's *t* test (*P<0.05; ***P<0.001 as compared with cyclophosphamide-treated group at 100 mg/kg).

### Materials and Methods

**Animals and injection**. All procedures involving mice and care have been conducted at the Interdepartmental Service Centre- Station for Animal Technology (STA), University of Rome "Tor Vergata", in accordance with the ethical standards, according to the Declaration of Helsinki, in compliance with our institutional animal care guidelines and following national and international directives (Italian Legislative Decree 26/2014, Directive 2010/62/EU of the European Parliament and of the Council). The ovaries were collected from CD-1 mice (Charles River) of 6 to 8 days old. Newborn mice (P6) were treated with intraperitoneal (ip) injection with PBS or cyclophosphamide (50, 75, 100 and 200 mg per kg of body weight). Mice were pre-treated with different inhibitors for 1 h before cyclophosphamide injection using a sterile micro syringe (Becton Dickson). Inhibitors used: GNF2 (range 15 mg -26 mg per kg of body weight), IMATINIB (12.3 - 41.1 mg per kg of body weight), NU (1.8- 5.3 mg per kg of body weight), KU (10 mg per kg of body weight). Cyclophosphamide (BAXTER) was prepared fresh as concentrated 40mg/ml in PBS. We dissolved Imatinib methane sulfonate salt (Novartis) in water, GNF-2 (SIGMA), NU (TOCRIS) and KU55993 (TOCRIS) in DMSO.

**Immunohistochemistry, Follicle counting and statistical analysis.** We prepared sections from ovaries fixed in MetaCarnoy solution (as previously described (Gonfloni et al 2009), embedded in paraffin and cut in slices of 5-7 µm of thickness. Sections were dewaxed, re-hydrated, and microwaved. Slices were then permeabilized with PBS triton 0,2 % and incubated with MSY-2 antibody (Santa Cruz). The staining was performed with immunocruz staining system for anti-goat antibody (Santa Cruz, sc-2023) and 3-aminoethyl-9-ethylcarbazole as substrate (AEC, Sigma). Sections were counterstained with hematoxylin and cover-slipped with Aquatex. Quantification of primordial and primary or secondary follicles was derived from histological analysis, counting Msy2-positive germ cells of mid-ovary sections. For each ovary (P9), several central slices (10<n<15) are included in the counting, with the exception of smaller peripheral slices (12-14 on average per each ovary). Quantification of primordial/primary follicle reserve is expressed as mean of immature follicles (primordial plus primary follicles) per single ovary. Average values for each ovary are represented as discrete points on a scatter plot. Mean value ± S.D. are shown in the scatter plot. The analysis of variance is evaluated with one-way ANOVA, with Turkey multiple comparison Test using PRISM 6 (Graph Pad software) (*$P$<0.05; **$P$<0.01; ***$P$<0.001) or by unpaired Student's *t* test where indicated.

**Immunofluorescence.** We prepared sections from ovaries sections fixed in MetaCarnoy solution, embedded in paraffin and cut in slice of 5-7 μm of thickness. Sections were dewaxed re-hydrated and microwaved in sodium citrate 10mM pH6, to expose the antigens. Unspecific-binding sites were blocked by incubating sections for 2hrs in a blocking solution (PBS plus 1%glycine, 5% BSA, 5% FBS and 5% NGS (normal goat serum). Ovaries sections were then incubated overnight with antibodies against p63, MSY-2 p-ATM, γH2AX, p-DNA-PK, p-p53, p-CHK2, p-DNAPK, pAKT, p-FOXO3a and cleaved PARP. After washing in PBS triton 0,05%, tissue sections were incubated with Alexa 555-goat anti-mouse (life technologies) and alexa 488-goat anti rabbit (invitrogen).

**Immunoblot analysis**. P7 dry ice–frozen ovaries were homogenized with a mini-pestle in ice-cold lysis buffer (50 mM Tris-HCl pH 7.5, 150 mM NaCl, 0.5% NP-40, 5 mM EDTA, 0.5% sodium deoxycholate, 1 mM phenylmethylsulfonyl fluoride, 1 mM sodium o-vanadate, 10 μg $ml^{-1}$ Tosyl phenylalanyl chloromethyl ketone (TPCK), 10 μg $ml^{-1}$, Tosyl-L-lysyl-chloromethane hydrochloride (TLCK) supplemented with protease inhibitors, all purchased from SIGMA). Equal amounts of protein extract (equivalent of one up to three ovaries) was loaded onto 6%, 8% or 12% SDS-PAGE gel and transferred to a nitrocellulose membrane (Amersham Bioscience).

**Tunel.** Ovary sections were stained according to the Fluorescein In Situ Cell Death detection Kit (Roche Diagnostic) and analyzed with a fluorescent filter. We used the protocol recommended by the manufacturer. DAPI (Molecular Probes Inc.)

**Reagents**. Antibodies Msy-2, Abl K-12, β-tubulin and were purchased from Santa Cruz; antibody for p-H2AX and H2AX and p-Tyr (4G10) were purchased from Millipore; antibody for p-DNA-PK (S2056), antibody for P63 (Y4A3) were purchased from SIGMA, antibody for p-ATM (S1981) was purchased from Rockland, polyclonal antibody for p63 was an home-made rabbit serum; antibody for p-P53 (S15), p-CHK2, p-FOXO3a and p-CHK2 were purchased from Cell Signaling Technology. Antibody for Abl (8E9) was purchased from BD-pharmingen. Secondary antibodies were purchased from Jackson Immunoresearch. All the antibodies were diluted in a blocking solution containing 5% BSA in PBS tween 0,05% for Western Blotting analysis and in a blocking solution containing 1% glycine, 5% FBS, 5% BSA and 5% NGS for immunofluorescence.

**Mating protocol.** We injected five cohorts of newborn CD1 mice (25 total female pups) with a single dose of cyclophosphamide (100 mg per kg of body weight), GNF-2 (26 mg per kg body weight) and with cyclophosphamide in combination with GNF2 or DPH (20 mg per kg body weight). Injection with PBS was used as a control. Five-six weeks after injection, we mated the control and treated mice with proven fertile males at regular time intervals every 5–6 weeks. Once mating was established by the formation of the fertilization plug (this occurs within a week), we separated the females and allowed the pregnancies to progress until delivery. We kept the mice for a week with their pups and then separated them. After a week (without breast-feeding), we mated them again with proven fertile males.
